# Cell-specific regulation of gene expression using splicing-dependent frameshifting

**DOI:** 10.1101/2022.03.02.481623

**Authors:** Jonathan P. Ling, Alexei M. Bygrave, Clayton P. Santiago, Rogger P. Carmen-Orozco, Vickie Trinh, Minzhong Yu, Yini Li, Jeong Han, Kamil Taneja, Ying Liu, Rochinelle Dongmo, Travis A. Babola, Patrick Parker, Lizhi Jiang, Patrick J. Leavey, Jennifer J. Smith, Rachel Vistein, Megan Y. Gimmen, Benjamin Dubner, Eric Helmenstine, Patric Teodorescu, Theodore Karantanos, Gabriel Ghiaur, Patrick O. Kanold, Dwight Bergles, Ben Langmead, Shuying Sun, Kristina J. Nielsen, Neal Peachey, Mandeep S. Singh, W. Brian Dalton, Fatemeh Rajaii, Richard L. Huganir, Seth Blackshaw

## Abstract

Precise and reliable cell-specific gene delivery remains technically challenging. Here we report a splicing-based approach for controlling gene expression whereby separate translational reading frames are coupled to the inclusion or exclusion of cell-specific alternative exons. Candidate exons are identified by analyzing thousands of publicly available RNA sequencing datasets and filtering by cell specificity, sequence conservation, and local intron length. This method, which we denote splicing-linked expression design (SLED), can be combined in a Boolean manner with existing techniques such as minipromoters and viral capsids. SLED vectors can leverage the strong expression of constitutive promoters, without sacrificing precision, by decoupling the tradeoff between promoter strength and selectivity. We generated SLED vectors to selectively target all neurons, photoreceptors, or excitatory neurons, and demonstrated that specificity was retained *in vivo* when delivered using AAVs. We further demonstrated the utility of SLED by creating what would otherwise be unobtainable research tools, specifically a GluA2 flip/flop reporter and a dual excitatory/inhibitory neuronal calcium indicator. Finally, we show the translational potential of SLED by rescuing photoreceptor degeneration in *Prph2*^*rds/rds*^ mice and by developing an oncolytic vector that can selectively induce apoptosis in SF3B1 mutant cancer cells. The flexibility of SLED technology enables new avenues for basic and translational research.

## Introduction

Cell type-specific control of gene expression is essential for both basic and translational biological research. Though this is often achieved using transgenic animal models, these are costly, difficult to scale, and restricted to a limited number of model organisms. An alternative approach, which is directly applicable for therapeutic purposes, is to use exogenous viral or plasmid constructs to selectively express genes of interest in specific cell types ^1–3^. These methods rely on the use of minimal promoters and enhancers that place constructs under the regulation of cell type-specific transcription factors ^4–8^, unique capsid proteins or surface features to limit the range of cell types infected by viral constructs ^9,10^, or the inclusion of specific miRNA seed sequences to inhibit off-target expression ^11–13^.

Current approaches, however, have important limitations. Minipromoter and enhancer-based constructs are difficult to develop and test in a systematic manner. For example, when removed from their genomic context, or tested in other species, they often show unpredictable patterns of cell-specific expression, despite showing high sequence conservation and patterns of chromatin accessibility ^14^. Furthermore, while viral serotypes typically provide enriched cell-specificity, thus far they are not strictly cell type-specific, and are not relevant for viral-independent gene delivery strategies. Likewise, microRNA-based approaches can help reduce off-target delivery in certain cells, but must be used in conjunction with other methods to achieve cell type-specific expression.

An orthogonal strategy that can be combined with the above approaches would be to harness alternative splicing of mRNA (RNA splicing) to direct cell-specific gene expression. RNA splicing is a highly regulated process that generates transcriptomic and proteomic diversity and many splicing patterns are correlated with unique cell types or cellular states. Fluorescent reporter vectors have been used to study the mechanistic regulation of alternative splicing events ^15–18^, but the large size of most intronic sequences precludes their inclusion in the most commonly used viral vectors. Adeno-associated virus (AAV) vectors are a leading platform for gene therapy due to their demonstrated safety and long-term efficacy across a variety of tissues ^19–22^, but these viruses are limited by a maximum packaging size of ∼4.7kb ^23^. Since the average intron length in the human genome is ∼5.4kb in length ^24^, it has been historically difficult to identify cell type-specific patterns of alternative splicing that are potentially compatible with AAV vectors ^25,26^. However, rapid adoption of full-length RNA sequencing (RNA-Seq) over the past decade has led to the public archival of datasets obtained from various cell types across multiple species. Furthermore, recent computational methods have been developed to comprehensively analyze patterns of alternative splicing across hundreds of thousands of publicly archived RNA-Seq datasets ^27–29^. We have used these databases to identify many cell type-specific alternative exons that are suitable for use in AAV vectors.

In this study, we have developed a suite of AAV-based tools that direct pan-neuronal, excitatory neuron, and photoreceptor-specific gene expression via a <u>s</u>plicing-linked <u>e</u>xpression <u>d</u>esign (SLED) strategy. This method uses splicing-dependent frameshifting, in combination with both ubiquitous and cell type-specific promoters, to drive cell type-specific expression of fluorescent proteins and other genes of interest. We show that, due to their small size, SLED constructs can be packaged into AAV vectors and that cell specificity is maintained *in vivo* across multiple species. Furthermore, the SLED method can be used to create previously unobtainable research tools. We miniaturized the *Gria2* flip/flop intron for AAV packaging to monitor this mutually exclusive splicing event at single-neuron resolution. We also demonstrated that dual calcium sensors can be simultaneously expressed in different cell types using a single expression vector, instead of using multiple viruses or transgenic animals. Finally, we demonstrated that SLED-based AAV constructs perform as efficiently as state-of-the-art minipromoter vectors for functional rescue of photoreceptor dystrophies, and also show that SLED can be used to selectively target *SF3B1* mutant cancer cells for oncolytic therapy. These results demonstrate that SLED-based tools are compatible with existing methods for regulating cell type-specific gene expression, and that SLED is broadly useful for a range of basic and translational research applications.

## Results

### Identification of cell-specific exons and SLED vector construction

To test the ability of alternative splicing to mediate cell type-specific expression of reporter constructs, we modified an existing bichromatic reporter plasmid ^15^. In this construct, dsRed is expressed when the default splicing pathway is used. When a cell type-specific alternative exon is spliced in, however, this results in a reading frame shift that leads to the expression of EGFP. In cases where the sequence length of the cell type-specific exon is a multiple of 3 and lacks a stop codon in the initiating translational reading frame, point mutations were introduced to create a frameshifting cell type-specific exon. Importantly, the dsRed sequence is modified to remove stop codons that would otherwise occur in the EGFP reading frame (Fig. 1a). 2A self-cleaving peptide sequences ^30^ were also included in front of each fluorescent protein to allow expression of the fluorescent protein independent of leader sequences (Fig. S1).

**Figure 1.**
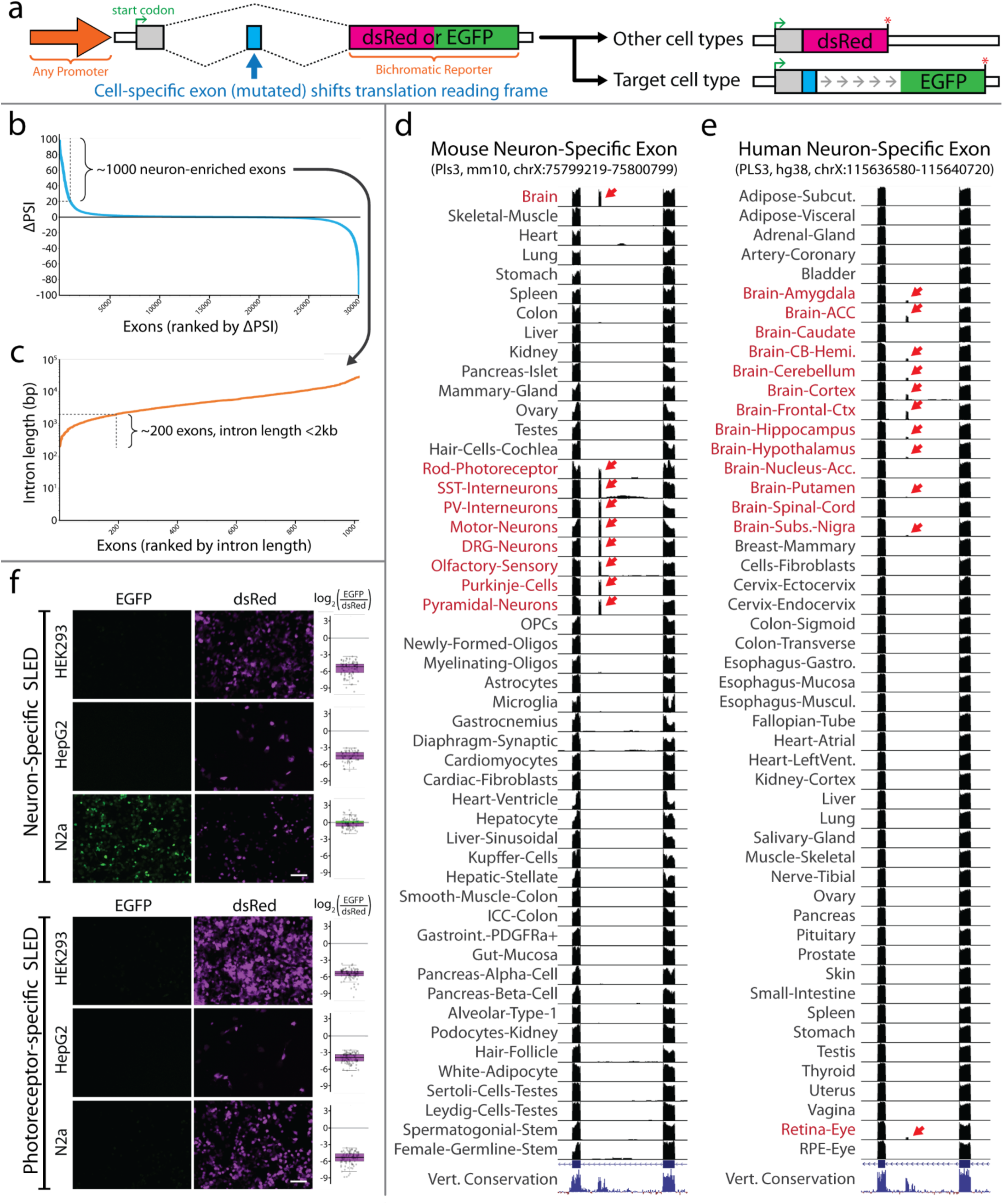
Identification of cell-specific exons and SLED vector construction. (**a**) Diagramatic sketch of SLED vector design strategy. SLED is compatible with any promoter. A frameshifting mutation is introduced into a cell-specific alternative exon to create two potential translational reading frames from an upstream start codon. In most SLED vectors, exon skipping will produce a red fluorescent protein while exon inclusion will shift the reading frame to produce a green fluorescent protein. (**b**) Ranking all neuron-enriched exons by the percent spliced-in (PSI) difference between neurons and other cell types. Exons were identified from mouse RNA-Seq datasets analyzed with the ASCOT pipeline ^27^. Approximately 1000 neuron-enriched exons have a ΔPSI greater than 20. (**c**) Among these top 1000 exons, approximately 200 candidates reside in introns <2kb in length. (**d, e**) UCSC genome browser views of the neuron-specific exon in *Pls3* that is used in SLED.NPL. Exon incorporation is only observed in neuronal datasets (red arrows) from both mouse (**d**) and human (**e**). A similar strategy was used to identify the photoreceptor-specific exon in *Atp1b2*. These exons were used to generate SLED vectors that were then tested in HEK293, HepG2, and N2a cancer cell lines (**f**). As predicted, neither vector showed EGFP expression, indicating an absence of cell-specific exon incorporation, except when the neuron-specific SLED was transfected into N2a cells, reflecting the neuronal characteristics of N2a neuroblastoma cells. Scale bars = 50μm.

We used three criteria to select alternative exons for analysis. First, alternative exons needed to show highly cell type-specific patterns of inclusion. Second, cell type-specific patterns of splicing needed to be conserved between mice and humans. Finally, the size of the intronic sequence used needed to be less than 2 kb. Using a computational resource that catalogs cell type-specific splicing patterns (ASCOT) ^27^, we identified ∼1000 neuronal-enriched alternative exons, of which ∼200 had intronic lengths of <2 kb (Fig. 1b,c). ∼99% of exons show high conservation (vertebrate phyloP score >1.5) of neuron-enriched splicing between mouse and human (Fig. 1d,e). A neuronally-enriched exon in the gene encoding the ubiquitously-expressed actin-binding protein Plastin 3 (*PLS3*) was selected for characterization. A similar process was used to identify a photoreceptor-specific exon in the gene encoding the ubiquitously expressed subunit of the ATPase Na+/K+ Transporting Subunit Beta 2 (*ATP1B2*) (Fig. S2). For proof-of-concept, we transfected the pan-neuronal and photoreceptor-specific SLED vectors into HEK293, HepG2, and N2a neuroblastoma cell lines to determine specificity (Fig. 1f). While dsRed was expressed in all cells, EGFP was only observed with pan-neuronal SLED in N2a cells, which exhibit neuronal precursor-like characteristics ^31^. No expression of EGFP was observed in any cells when transfected with the photoreceptor-specific SLED construct, supporting the cell type specificity of the *ATP1B2* alternative exon. Specificity was determined at the single cell level by evaluating the log_2_ ratio of EGFP/dsRed fluorescence (Fig. 1f).

### SLED vectors maintain cell-specific expression when delivered using AAV

To test whether SLED vectors retain specificity when packaged into AAV, we cloned pan-neuronal SLED (SLED.NPL) and photoreceptor-specific SLED (SLED.RAB) into an AAV backbone (Fig. 2a). Furthermore, we sought to test whether cell type-specific minipromoters could be combined with SLED-based constructs in a Boolean manner to provide more selective cell type-specific expression using AAV vectors. To do this, we combined the pan-neuronal hSyn minipromoter ^32^ with an alternative exon of the gene that encodes the ubiquitously expressed clathrin complex interactor Synergin gamma (*SYNRG*), which in the brain is specific to excitatory neurons and glia (SLED.ENS, Fig. 2a, Fig. S2).

**Figure 2.**
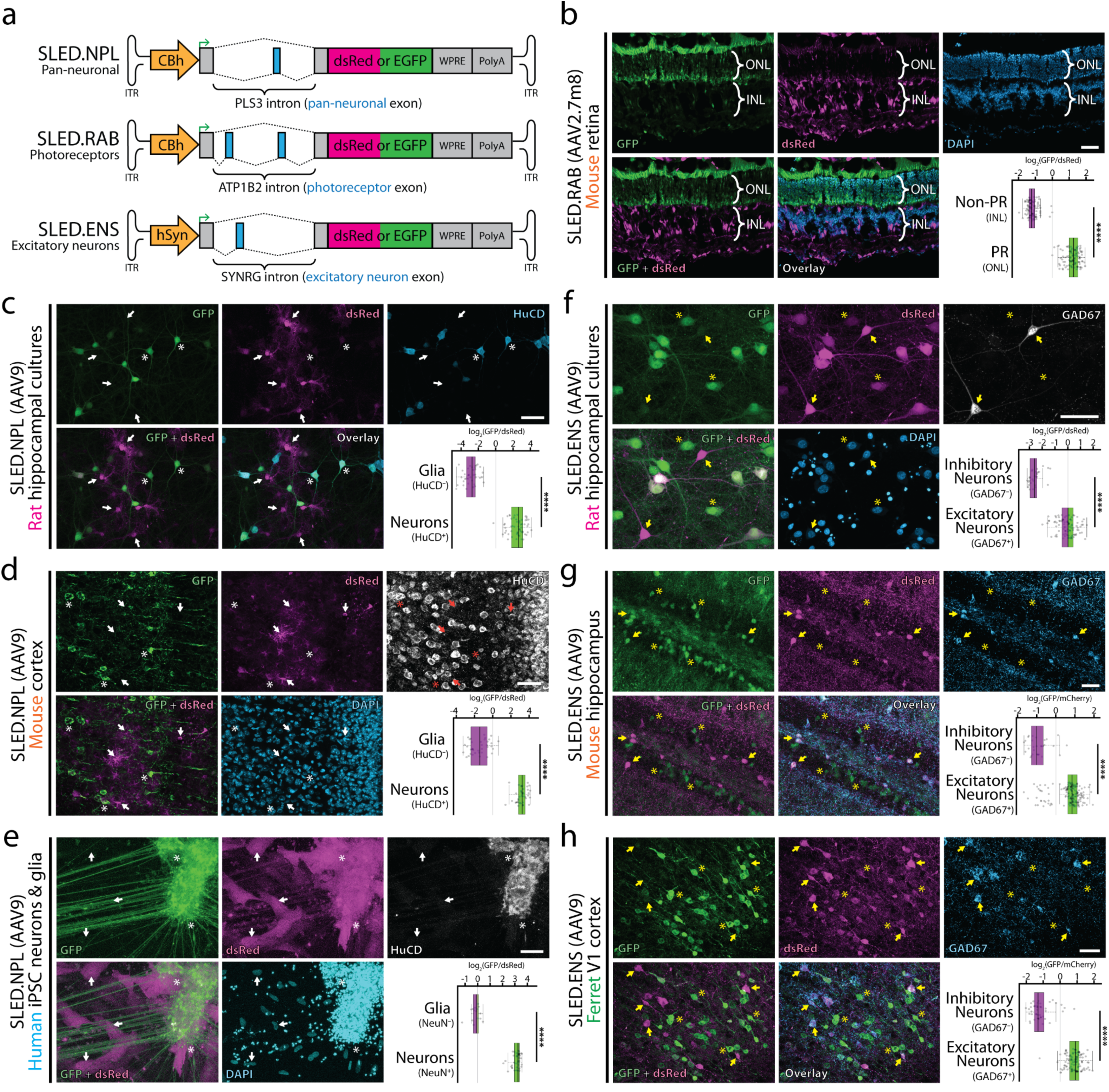
SLED vectors maintain cell-specific expression when delivered using AAV. (**a**) Diagramatic sketch of pan-neuronal (SLED.NPL), photoreceptor-specific (SLED.RAB), and excitatory neuron-specific (SLED.ENS) vectors designed for AAV packaging. In all SLED vectors, EGFP is translated when cell-specific exons are spliced-in. (**b**) SLED.RAB, packaged in AAV2.7m8, was intravitreally injected into P0 mouse retinas and processed at P30. EGFP is highly enriched in retinal photoreceptors while inner retinal neurons are strongly positive for dsRed. ONL = outer nuclear layer, INL = inner nuclear layer. (**c**-**e**) SLED.NPL, packaged in AAV9, was used to transduce primary rat hippocampal cultures (**c**), mouse cortex (**d**), and human iPSC-derived neurons (**e**). In all cases, EGFP is highly enriched in HuC/D^+^ (rat, mouse) or NeuN^+^ (human) neurons, while non-neuronal cells are strongly positive for dsRed. (**f**-**h**) SLED.ENS, packaged in AAV9, was used to transduce primary rat hippocampal cultures (**f**), mouse hippocampus (**g**), and ferret cortex (**h**). In all cases, EGFP is highly enriched in GAD67^−^ excitatory neurons, while GAD67^+^ inhibitory neurons are strongly positive for dsRed. **** indicate p < 0.0001, two-tailed t-test. For ratio calculations in panels b to h, n=205 (b), n=94 (c), n=78 (d), n=55 (e), n=121 (f), n=198 (g), and n=154 (h). Scale bars = 50μm.

We first tested the expression of these constructs using plasmid transfection and electroporation. As expected, we observed selective expression of EGFP in neurons following transfection of SLED.NPL into primary rat hippocampal cultures, although expression of the default splicing-driven dsRed was observed in transfected neurons and glia (Fig. S3). Likewise, in neonatal mouse retinal explants electroporated with the SLED.RAB construct ^33^, we observed expression of dsRed in all postnatally-generated cell types, but EGFP reporter expression is restricted to photoreceptors (Fig. S3). Lastly, we observe that transfection of the SLED.ENS constructs into primary rat hippocampal cultures resulted in selective exclusion of hSyn-driven EGFP expression from somatostatin-expressing GABAergic interneurons (Fig. S3). To further validate the photoreceptor-specificity of the SLED.RAB construct *in vivo*, postnatal day 0 (P0) mouse retinas were transduced with photoreceptor-specific AAV2.7m8.SLED.RAB and processed 4 weeks later at P30. This revealed highly enriched expression of EGFP in retinal photoreceptors, with inner retinal neurons strongly positive for dsRed (Fig. 2b).

We next tested the neuronal-specificity of SLED.NPL packaged into AAV9 by transducing primary rat hippocampal cultures at 1 day *in vitro* (DIV). At DIV 15, cells were fixed and immunofluorescence conducted for the neuronal marker HuC/D.. Comparison of the ratio of EGFP to dsRed fluorescence revealed that EGFP was highly enriched in HuC/D-positive neurons (Fig. 2c). To test SLED.NPL *in vivo*, we performed stereotactic injection of AAV9.SLED.NPL into the mouse hippocampal region. Efficient and widespread infection was observed and the neuronal specificity of EGFP expression was maintained (Fig. 2d). AAV-based gene therapies are being explored as treatment options for neurological disorders and SLED vectors may improve the safety and efficacy of these methods. To determine whether AAV9.SLED.NPL maintains neuron-specific expression in human cells, we transduced human iPSC-derived mixed neuronal and glial cultures (Fig. 2e). Here too, we observed strong EGFP expression in NeuN-positive neurons but only dsRed expression in NeuN-negative glial cells.

Lastly, we tested the specificity of the excitatory neuron-specific AAV9.SLED.ENS construct. Primary rat hippocampal cultures show strong expression of EGFP in Gad67-negative excitatory neurons, but little or no expression of EGFP in Gad67-positive GABAergic interneurons (Fig. 2f). Stereotactic injection of AAV9.SLED.ENS into mouse hippocampus likewise resulted in strong and broad neuronal expression of EGFP, but exclusion of EGFP signal from dsRed-positive, Gad67-positive interneurons (Fig. 2g). Finally, a similar pattern of exclusion from Gad67-positive interneurons was observed following transduction of primary ferret visual cortex (Fig. 2h). Together, these findings demonstrate that SLED cell specificity is maintained *in vitro* and *in vivo* across multiple species, and that mutually exclusive splicing events can be simultaneously monitored using AAV-based SLED tools.

### Generation of unique splicing-based tools using SLED vectors

We next sought to test whether SLED-based AAV vectors could be adapted to study *in vivo* patterns of flip/flop splicing in the AMPA-type glutamate receptor subunit GluA2. Flip/flop alternative splicing occurs within the ligand binding domain, and influences AMPA receptors assembly and channel kinetics ^34–37^. The short lengths and high sequence similarity of the mutually exclusive flip/flop exons precludes the use of immunostaining or *in situ* hybridization to detect their localization *in situ*, which has effectively restricted previous efforts investigating flip/flop splicing in learning and plasticity to using qRT-PCR analysis ^38,39^. To address this, we generated an hSyn-driven AAV vector which expresses EGFP when the flop exon is incorporated, and dsRed when the flip exon is incorporated (SLED.GluA2, Fig. 3a). To validate that AAV9.SLED.GluA2 splicing reflected endogenous flip/flop splicing, we electroporated primary rat cortical neurons and FACS-isolated high EGFP-expressing cells. We designed primers to selectively amplify spliced flip/flop exons in GluA2 mRNA (Fig. 3b). Digestion using HpaI, which selectively cleaves the flop exon into two smaller fragments, revealed expected enrichment of flop exon fragments in the EGFP-enriched fraction (Fig. 3c). This was further confirmed using Sanger sequencing of amplified products (Fig. 3d, Fig. S4).

**Figure 3:**
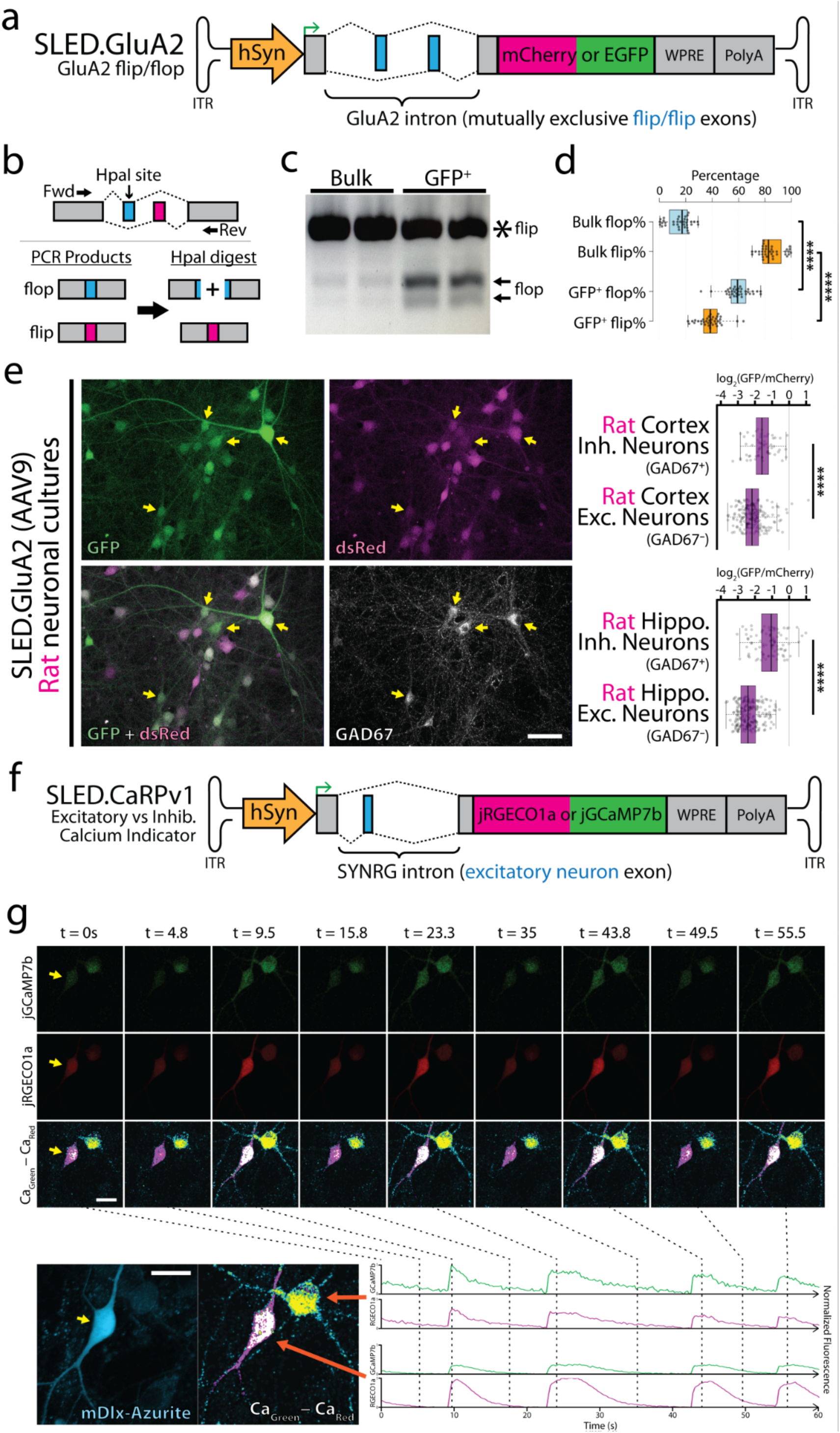
Generation of unique splicing-based tools using SLED vectors. (**a**) Diagramatic sketch of GluA2 (*Gria2*) flip/flop SLED vector design (SLED.GluA2). (**b**) To validate that SLED.GluA2 reflected endogenous GluA2 flip/flop splicing patterns, we designed endogenous mRNA-specific primers to PCR amplify the GluA2 flip/flop locus. Although the mutually exclusive flip and flop exons are identical in length and highly similar in sequence, HpaI will selectively digest the flop PCR product into two fragments. SLED.GluA2, packaged into AAV9, was used to electroporate primary rat neuronal cultures and EGFP^high^/mCherry^low^ cells were isolated using FACS (Fig. S4). RNA was extracted from EGFP^high^/mCherry^low^ cells and bulk rat neuronal cultures and primers (**b**) were used to amplify PCR products. (**c**) HpaI incubation yielded digestion products in the EGFP^high^/mCherry^low^ cells, which was further confirmed using Sanger sequencing (**d**, Fig. S4, n=36, **** indicate p < 0.0001, two-tailed t-test.). (**e**) Primary rat neuronal cultures transduced with SLED.GluA2 show significantly different EGFP/mCherry ratios between excitatory (GAD67^−^) and inhibitory (GAD67^+^) neurons. **** indicate p < 0.0001, two-tailed t-test. (**f**) Diagramatic sketch of bicistronic jRGECO1a (inhibitory neurons) and jGCaMP7b (excitatory neurons) SLED vector design (SLED.CaRPv1). (**g**) Transfection of primary rat neuronal cultures yielded divergent ratios in jGCaMP7b and RGECO1a intensities in excitatory (mDlx-Azurite^−^) and inhibitory (mDlx-Azurite^+^) neurons. Data presented represent jGCaMP7b (top row) and jRGECO1a (middle row) intensity values over a 60s time-lapse (4Hz). Bottom row represents a normalized representation of total Ca intensity scaled by the delta between jGCaMP7b and jRGECO1a pixel values. Individual Ca indicator traces are demonstrated in the bottom panels. For ratio calculations in panel e, n=266 (cortex) and n=316 (hippocampus). Scale bars = 50μm (panel e) and 20μm (panel g).

Transduction of AAV9.SLED.GluA2 into both primary rat hippocampal and cortical cultures (Fig. 3e) revealed a variety of different cellular patterns of reporter expression, with EGFP-dominant, dsRed-dominant and mixed cells all present. However, we observe that Gad67-positive hippocampal neurons are enriched for EGFP-dominant expression, matching previous observations obtained using single-cell SMART-Seq analysis and bulk RNA-Seq analysis of RiboTRAP-expressing interneurons ^40–42^ (Fig. S5).

The translational frameshifting used in SLED vectors also offers the potential to deliver multiple functional payloads, such as genetically encoded calcium sensors or optogenetic actuators, using a single viral vector. As proof of concept, we created a bicistronic calcium indicator vector based on the excitatory neuron-specific SLED.ENS. We termed this calcium reporter plasmid version 1 (CaRPv1) (Fig. 3f). In CaRPv1, GCaMP7b is expressed in excitatory neurons, while RGECO1a is expressed in the default translational reading frame (inhibitory neurons) ^43,44^. Identification of excitatory vs inhibitory neurons was established using the Δpixel intensity of normalized fluorescence values, due to differences in dynamic range and baseline fluorescence at resting calcium concentrations for GCaMP7b and RGECO1a (see methods). Transfection of CaRPv1 into primary rat hippocampal cultures revealed the expected patterns of calcium transients (Fig. S6, Supplemental Videos 1 & 2). Furthermore, large and synchronous calcium transients were observed following the addition of bicuculline (a GABA_A_ receptor antagonist, used to induce disinhibition), indicating that CaRPv1 was reporting cellular activity as expected (Fig. 3g). In its current design, CaRPv1 is unable to be packaged into AAV due to the size of the ENS intron (∼1600bp). However, future deletion mutagenesis and sequence optimization should enable AAV packaging of CaRPv1.

### Adapting SLED vectors for translational studies

Mutations in the photoreceptor outer segment structural gene *PRPH2* cause human retinal degeneration ^45^ and null mutations in *Prph2* lead to slow-onset photoreceptor degeneration in mice ^46–49^. AAV-based constructs driven by the photoreceptor-specific minipromoter mOps have been previously used to rescue *Prph2* expression in *rds/rds Prph2*-deficient mice (Prph2^*rds/rds*^), although only modest photoreceptor preservation, and no long-term recovery of visual function was observed due to weak promoter efficiency ^50^. We modified the photoreceptor-specific SLED.RAB to selectively express *PRPH2* under control of the ubiquitous CBh promoter. In parallel, we generated mOps minipromoter-driven rescue vectors that were used in previous studies ^50–52^. CMV-driven EGFP vectors were also obtained as controls (Fig. 4a). These were all packaged into AAV2.7m8 capsids and injected subretinally at P28 into Prph2^*rds/rds*^ mice.

**Figure 4:**
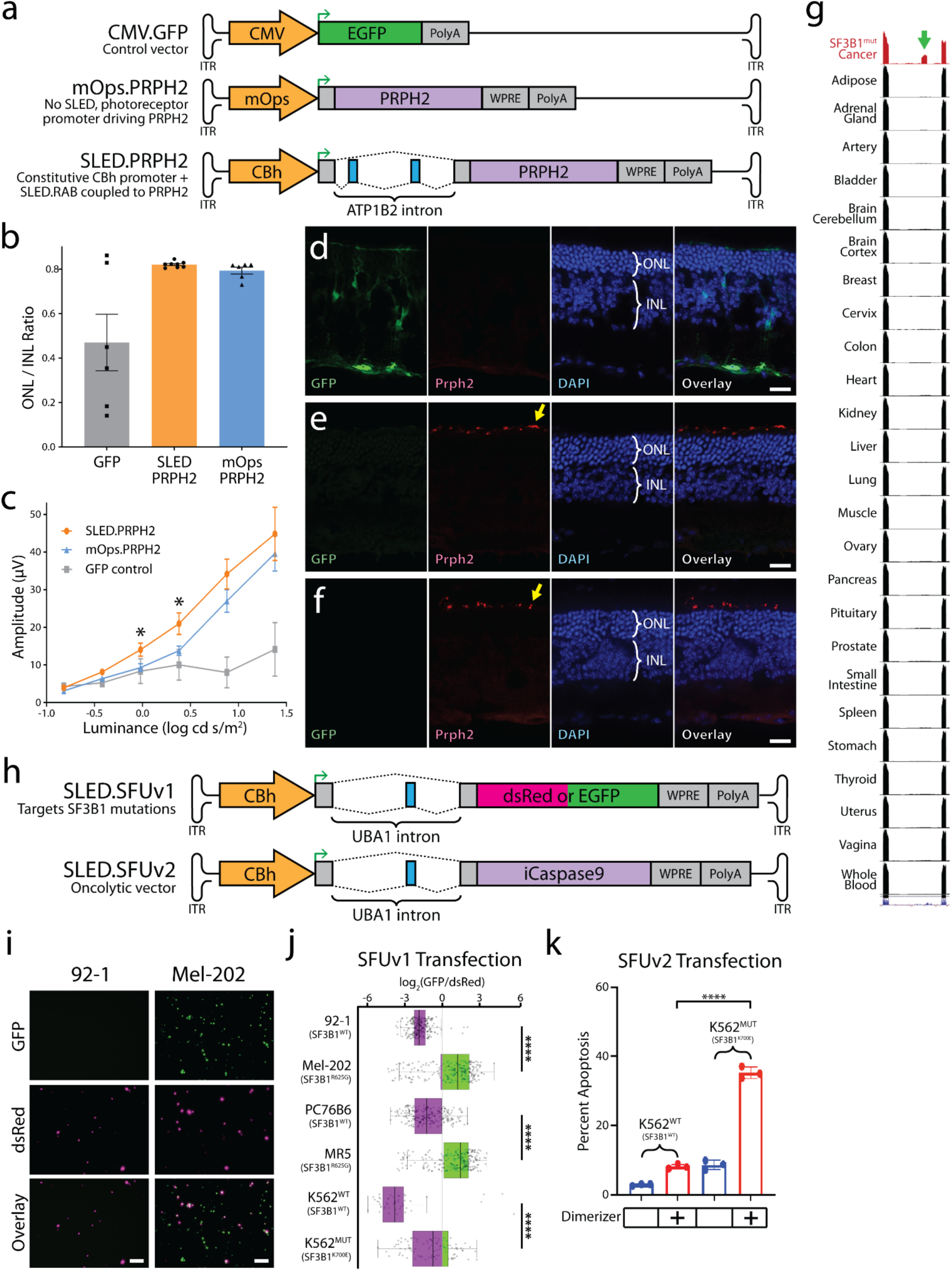
Adapting SLED vectors for translational studies. (**a**) Diagramatic sketch of GFP under the control of the constitutive CMV promoter (CMV.GFP), PRPH2 under the control of the photoreceptor specific mOps promoter (mOps.PRPH2), and PRPH2 regulated by the photoreceptor-specific SLED.RAB (SLED.PRPH2) vectors designed for AAV packaging. CMV.GFP, mOps.PRPH2, and SLED.PRPH2 were packaged into AAV2.7m8 for testing in *Prph2*^*rds/rds*^ animals. For experimental design, n=6 for each AAV treatment. (**b**) Average ONL/INL ratios and (**c**) average light-adapted ERG b-wave amplitudes in three month post-injected Prph2^*rds/rds*^ animals treated with CMV.GFP, SLED.PRPH2 or mOps.PRPH2 viral constructs (asterisks indicate p < 0.05, two-tailed t-test, comparison between mOps.PRPH2 and SLED.PRPH2). (**d**-**f**) Immunofluorescence staining of retinal sections from *Prph2*^*rds/rds*^ eyes injected with CMV.GFP (**d**), mOps.PRPH2 (**e**), and SLED.PRPH2 (**f**). ONL = outer nuclear layer, INL = inner nuclear layer. EGFP is only detected in CMV.GFP treated controls, but Prph2 signal (yellow arrow) is detected in photoreceptor inner segments of retinas transduced with both mOps.PRPH2 and SLED.PRPH2. (**g**) UCSC genome browser view of a cryptic exon in *UBA1* (green arrow) that is present in cancers with oncogenic SF3B1 mutations (TCGA ^76^) and absent in all normal human tissues sequenced by the GTEx consortium ^77^. (**h**) Diagramatic sketch of bichromatic fluorescent reporter based on the SF3B1^mut^-associated exon (SLED.SFUv1) and a similar vector where an inducible iCaspase9 kill switch is coupled to incorporation of the SF3B1^mut^-associated exon. (**i**) As a proof of concept, SLED.SFUv1 was transfected into uveal melanoma cell lines with (Mel-202) and without (92-1) *SF3B1* mutations. EGFP was highly enriched in only Mel-202 cells while dsRed was strongly expressed in 92-1 cells, which was validated using FACS (**j**). Isogenic cell lines derived from Mel-202 with the *SF3B1*^*R625G*^ mutation genetically inactivated (PC76B6) and maintained (MR5) showed similarly concordance, with strong EGFP expression only present in the *SF3B1*^*R625G*^ MR5 cell line. Likewise, strong EGFP expression was only present in *SF3B1*^*K700E*^ K562 leukemia cells, as compared to wildtype K562 cells. **** indicate p < 0.0001, two-tailed t-test. For ratio calculations, n=245 (92-1), n=1158 (Mel-202), n=377 (PC76B6), n=526 (MR5), n=49 (K562^WT^), n=677 (K562^MUT^). (**k**) Transfection of wildtype K562 cells and mutant *SF3B1*^*K700E*^ K562 cells with SLED.SFUv2 revealed strong apoptosis only in *SF3B1*^*K700E*^ K562 cells treated with the iCaspase9 activating dimerizer (n=3 FACS replicates). Scale bars = 50μm.

Mice were injected with AAV at one month of age, and three months later were then analyzed using optical coherence tomography (OCT) to measure the relative thickness of the retinal outer nuclear layer (ONL), where photoreceptors reside. We observed that ONL thickness was similar in both SLED and mOps-regulated AAV constructs, and significantly greater than mice injected with CMV.GFP control virus (Fig. 4b). The amplitude of the light-adapted, cone-mediated, full-field electroretinogram (ERG) was larger in SLED relative to mOps-based rescue constructs, with both showing significantly higher b-wave responses relative to CMV.GFP controls (Fig. 4c). Immunostaining for Prph2 in transduced retina showed no detectable expression in CMV.GFP controls (Fig. 4d), but Prph2 signal was detected in photoreceptor inner segments in retinas transduced with both mOps (Fig. 4e) and SLED-based (Fig. 4f) Prph2 rescue constructs.

Finally, because oncolytic virotherapy is now an approved treatment modality in oncology ^53^, we sought to leverage tumor-specific RNA splicing patterns to generate SLED-based oncolytic vectors. Specifically, we identified a cryptic alternative exon in the constitutively expressed ubiquitin-like modifier activating enzyme 1 (*UBA1*) that was observed exclusively in *SF3B1* mutant cancer cells (Fig. 4g). We designed two SLED-based vectors incorporating this alternative exon: one that express a bichromatic fluorescent reporter that selectively expresses EGFP in *SF3B1* mutant cells (SLED.SFUv1, where EGFP will only express in mutant *SF3B1* cells), and one that selectively expresses an oncolytic inducible Caspase 9 ^54,55^ in *SF3B1* mutant cells (SLED.SFUv2) (Fig. 4h). We first tested SFUv1 specificity by transfected 92-1 and Mel-202 uveal melanoma cell lines. We observed that EGFP expression was present in *SF3B1*^R625G^ Mel-202 cell lines ^56^, but absent in the 92-1 uveal melanoma cell line, which is wildtype for *SF3B1* (Fig. 4i) ^57^. We next quantified this using FACS analysis and analyzed four additional cell lines, two of which were isogenic to Mel-202: PC76B6, in which AAV-based gene targeting was used to revert the mutant *SF3B1* status to wildtype through inactivation of the *SF3B1*^R625G^ allele, and MR5, a gene targeting control clone of Mel-202 that retains the *SF3B1*^R625G^ mutation (Fig. S7). The other two cell lines analyzed by FACS were wildtype and *SF3B1*^K700E^ K562 leukemia cells. FACS analysis revealed that the EGFP/dsRed ratio is strongly dependent on the presence of either the *SF3B1*^R625G^ or *SF3B1*^K700E^ mutation (Fig. 4j). Finally, we tested the efficacy of SLED.SFUv2 by transfecting the wildtype and *SF3B1*^K700E^ K562 cells and observed efficient and selective induction of apoptosis in cells carrying the *SF3B1*^K700E^ following induction of Caspase9 dimerization (Fig. 4k).

## Discussion

In this study, we demonstrate that SLED-based vectors can produce cell-specific expression in a variety of constructs, and that SLED-based approaches are both orthogonal and complementary to existing methodology. SLED-based alternative splicing can be combined in a Boolean fashion with minipromoters to achieve higher levels of cell type specificity, as demonstrated by the integration of the pan-neuronal hSyn minipromoter and the excitatory neuron and glial-specific *SYNRG* exon to generate the excitatory neuron-specific SLED.ENS vector. SLED-based AAV vectors can also be used to study previously intractable problems without the use of complex transgenics, such as the *in vivo* dynamics of GluA2 flip/flop splicing. The use of splicing-related frameshifting allows efficient cell type-specific expression of multiple reporter or effector constructs in a single vector. SLED-based vectors also enable new strategies to improve gene therapies. For instance, SLED vectors can use any promoter, potentially allowing for stronger and more sustained levels of expression relative to conventional minipromoters and enabling more consistent patterns of cell-specific expression across multiple model organisms ^58,59^. This is critical, as photoreceptor minipromoters have encountered complex issues when tested in various mammalian species. For instance, the hRK1 minipromoter, which is widely used to drive expression in both rods and cones in rodents, is unable to drive efficient expression in cones of other model organisms such as dogs and pigs unless used at very high titers, and expresses at lower levels in rods than the rod-specific mOps promoter ^59–61^. Finally, SLED vectors can also selectively target disease states associated with abnormal splicing that would not be accessible using minipromoters. The use of photoreceptor minipromoters in AAV vectors can lead to long-term toxicity, but this is avoided using constitutive promoters ^62^.

Identification of evolutionarily-conserved patterns of cell-specific alternative splicing is straightforward, provided that good quality full-length RNA-Seq data is available. As transcriptomes from more tissues and cell types are profiled and deposited in public archives, our ability to identify highly cell-specific patterns of alternative splicing will increase and these datasets will guide the design of the next generation of SLED vectors. While transcriptome analysis has increasingly shifted towards 3’-directed short read single cell RNA-Seq platforms in recent years, emerging techniques such as long-read nanopore sequencing ^63^ and economical full-length scRNA-Seq techniques such as SMART-Seq v3 will continue to improve our knowledge of splicing patterns ^64,65^. Recent compendia of splice-junction and transcript-level expression have surveyed 100,000s to millions of datasets ^28,29,66^, making these patterns easier to discover computationally.

With detailed characterization of alternative splicing patterns in the tremendous diversity of cell types, particularly in the human central nervous system, an intersectional approach combining SLED and cell-specific minipromoters may generate vectors that can selectively target to date untargetable cell types. Indeed, alternative splicing generates another layer of transcriptional complexity to the nervous system ^25,67,68^.

SLED-based vectors are still intrinsically limited by the size of the genomic intronic sequences used to control alternative splicing, which are generally substantially larger than minipromoters. While this is a less severe obstacle for transfection- or nanoparticle-based gene delivery, it is still a substantial limitation for AAV-based delivery. While the effects of deletion mutagenesis on cell-specific splicing can be unpredictable, recently developed machine learning algorithms may help facilitate rational design of smaller SLED vectors ^69,70^. Drug-inducible approaches to regulate splicing ^71–73^ and the inclusion of miRNA target sites ^74,75^ may enable further control of SLED-based constructs.

## Materials and Methods

### Molecular cloning and cancer cell line culture

To generate SLED plasmids, gene fragments were commercially synthesized using Twist Biosciences and ThermoFisher GeneArt and cloned into an AAV vector backbone (Addgene #105922) using restriction enzyme cloning. HEK293, HepG2, and N2a cells were cultured in Dulbecco’s Modified Eagle’s Medium (Corning, 10-017-CV) supplemented with 1x GlutaMAX (ThermoFisher Scientific, 35050061), 10% FBS (Corning, 35-010-CV). Human uveal melanoma cell lines 92-1 (generously provided by Charles Eberhart, Johns Hopkins University), MP41 (ATCC), and Mel-202 (Sigma) were cultured in RPMI medium with 10% fetal bovine serum (FBS), penicillin/streptomycin, and l-glutamine. SF3B1^K700E^ and control K562 cells were obtained from Horizon Discovery and cultured in RPMI with 20% FBS. The isolation, early characterization and further genetic and molecular characterization of the cell lines have been described elsewhere ^78–80^. Transfection of SLED vectors in uveal melanoma cells was achieved using Lipofectamine 3000 (ThermoFisher Scientific, L3000008) and with the 4D-Nucleofector X (Lonza) for K562 cells. The SF3B1^R702R^ AAV targeting vector as described ^81^ was applied to SF3B1^R625G^ Mel202 cells. iCaspase9 dimerization was induced by 100nM AP21087 (Sigma-Aldrich).

### Antibodies

The following antibodies were used for primary culture neurons (supplier, catalog number and working dilution are indicated): anti-Gad67 ms IgG2a (Millipore MAB5406, 1:500); anti-somatostatin rat (Millipore MAB354, 1:400); anti-HuCD IgG2B (Thermo 16A11, 1:200); anti-NeuN mouse IgG1 (Thermo MAB377, 1:500). The following antibodies were used for brain sections (supplier, catalog number and working dilution are indicated): anti-GFP chicken polyclonal (Abcam ab13970, 1:2000); anti-dsRED rabbit polyclonal (Tanaka LivingColors 632496, 1:1000) (this antibody also detects mCherry); anti-NeuN mouse monoclonal IgG1 (Thermo MAB377, 1:500); anti-Gad67 mouse monoclonal IgG2a (Millipore MAB5406, 1:200); anti-PV mouse monoclonal IgG1 (Swant PV235, 1:2000); anti-somatostatin rat monoclonal IgG2b (Millipore MAB354, 1:400).

### Preparation and treatment of rat primary cultures

Hippocampi and cortices were dissected from embryonic day 18 rats, incubated with papain (Worthington Biochemical) and gently triturated with polished glass pipettes. Hippocampal neurons were plated on 18mm glass coverslips precoated with poly-L-Lysine (1mg/ml) in NeuroBasal media (Gibco) supplemented with 2% B27 (Gibco), 50 U/ml penicillin, 50 mg/ml streptomycin, 2mM GlutaMax (Gibco) and 5% horse serum (Hyclone). Hippocampal cells were plated at a density of 150K/coverslip (in a 12-well plate). At days in vitro 1 (DIV1) media was replaced with NM0 consisting of the above plating media without the addition of serum. Cells were then fed every 7 days with NMO. Hippocampal cultures were transduced with viral vectors at DIV1 and fixed at DIV15 for immunofluorescence analysis. SLED constructs were also transfected into hippocampal neurons using Lipofectamine 2000 (Invitrogen) following the manufacturer’s instruction. Cortical neurons were used to evaluate SLED.GluA2. Before plating, cortical cells (6M per reaction) were electroporated with SLED.GluA2 plasmid DNA (3-5ug) using a Rat Neurofection kit (Amaxa) and split evenly between 3 wells of a 6-well plate. As a comparison group, neurons were plated at equivalent density without electroporation. Cortical cells were harvested at (DIV4-6) for FACS and downstream evaluation of Gria2 flip/flop splicing.

### Quantification of SLED.ENS with IMARIS

To calculate the green/red ratios of SLED.ENS in excitatory and inhibitory neurons, Z-stacks were imported into IMARIS (Bitplane version 9.7.0) and 3D surfaces created around the cell bodies of transduced cells. The average EGFP and mCherry signal within the 3D surface was extracted for each cell. The Gad67 immunohistochemical signal was used to classify the cells into Gad67 positive (inhibitory interneuron) or Gad67 negative (putative excitatory neurons). To determine the statistical specificity of SLED vectors, log base 2 transformed ratios of green/red fluorescence were compared between sample groups using unpaired t-tests (GraphPad).

### Quantification of SLED.ENS, SLED.NPL and SLED.GluA2 in Fiji

Confocal Z-stack images acquired with a 40x objective were processed using Fiji ^82^. Maximum intensity projections were generated, and the EGFP or mCherry/dsRed channel used to draw circle/oval ROIs around the cell body of transduced neurons without looking at signal in the 405 or 647 channels that contained cell-specific immunofluorescent markers. The raw EGFP and mCherry/dsRED average ROI intensities were then extracted for each cell. Subsequently, the cell-specific identification of each cell was observed from immunofluorescence using the 405 and 647 channel (for SLED.NPL this was HuC/D expression, for SLED.ENS this was Gad67 expression and for SLED.GluA2 this was Gad67). This enables clustering of the individual cells for comparisons of the green/red ratios. To determine the statistical specificity of SLED vectors, log base 2 transformed ratios of green/red fluorescence were compared between sample groups using unpaired t-tests (GraphPad).

### Live-cell imaging of SLED.CaRPv1

Rat primary hippocampal neurons at DIV11 were transfected with 1μg SLED.CaRPv1 alongside 1μg of mDlx-Azurite (as an enhancer based marker of interneurons ^4,83^) per coverslip in a 12-well plate. At DIV13 coverslips were imaged live in ACSF (NaCl 120mM, KCl 5mM, HEPES 10mM, D-Glucose 10mM, CaCl2.2H2O 2mM, MgCl2 1mM) at pH 7.4 using a Zeiss LSM 880 confocal microscope in a temperature (37°C) and humidity controlled chamber. Interneurons were identified by presence of the mDlx-Azurite signal, and mDlx-Azurite-negative cells were considered putative excitatory neurons. Time series were collected using 20x or 10x objectives for single cell or multiple cell imaging, respectively. Images were acquired at baseline, and also following addition of Bicuculline (20μM) to promote network activity through disinhibition. Files were processed and analyzed using Fiji ^82^. Fluorescence signals from jGCaMP7b and jRGECO1a were normalized to maximize variation between mDlx-Azurite positive and negative cells and the difference (Δintensity) between jGCaMP7b and jRGECO1a was calculated across each pixel and image frame (a gaussian blur (1px) was applied to each image before Δintensity to avoid pixelation artifacts). The sum of jGCaMP7b and jRGECO1a pixel values (sumGR) were calculated across each pixel and image frame. To generate the Ca_Green_-Ca_Red_ heatmap in Figure 3g and Supplemental Videos 1 and 2, sumGR was multiplied by Δintensity and colored using a custom lookup table.

### Immunofluorescence analysis of primary cultured neurons

Unless otherwise stated, hippocampal neurons were fixed at DIV15. Media was aspirated and cells were washed with PBS at RT once before being incubated with 4% PFA (Electron Microscopy Sciences) made up in PBS with the addition of 4% sucrose at RT for 15 mins. Coverslips were washed 4 times with PBS then immunofluorescence commenced using a gelatin-based buffer (15 mM phosphate buffer (pH 7.4) containing 0.1% gelatin, 0.3% Triton X-100, and 0.25 M NaCl) for combined blocking/antibody incubation. Primary antibodies (see below) were incubated with coverslips O/N at 4°C. Secondary antibodies (Invitrogen for 488, 568 and 647 and Jackson labs and Abcam for 405; all at 1:500 dilution) were incubated for 1hr at RT. Between antibody incubations cells were washed with PBS, with some experiments including a brief DAPI incubation to label cell nuclei. Coverslips were mounted on slides with PermaFluor (Thermo Fisher Scientific). Samples were imaged on a Zeiss LSM 880 confocal microscope. Care was taken to ensure pixels in each channel were not over saturated. Images were analyzed using Fiji. In cultured neurons the SLED-driven fluorophores were not antibody boosted.

### AAV production

SLED.ENS (1E+12 vg/ml, mouse and ferret cortex) and SLED.GluA2 (1E+12 vg/ml) were generated by the UNC Vector Core and SLED.NPL (2E+13 vg/ml) and SLED.RAB (2E+13 vg/ml) were generated by Virovek. SLED.ENS used for primary rat neuronal cultures was generated by Virovek (2E+13 vg/ml)

### Stereotaxic surgery and virus injections

All animals were treated in accordance with the Johns Hopkins University Animal Care and Use Committee (IACUC) guidelines. Adult Blk6/J mice were used to evaluate SLED.AAV vectors. Animals were anesthetized with isofluorane (Baxter) using a SomnoSuite system (Kent Scientific) and secured in a stereotaxic frame (Kopf). The animal’s temperature was controlled with a closed-loop system (RightTemp, Kent Scientific). The animal’s scalp skin was cleaned with an ethanol wipe, and the hair removed. Animals were injected with 0.5ml sterile saline (VetOne) to maintain hydration, buprenorphine (ZooPharm; 1 mg/ml) and lidocaine (VetOne; 2%) subcutaneously. The lidocaine was injected under the scalp as a local anesthetic. An incision was made to expose the skull surface, and to enable a small craniotomy to be made (see below for coordinates) exposing the brain surface. A glass pipette (Drummond Science Company; Wiretrol II) was pulled (Sutter Instruments) and sharpened to a 30° angle (Medical System Corp) and controlled by a pneumatic injector (Narishige) to enable controlled virus injection. Pipettes loaded with SLED viruses were slowly lowered to the desired stereotaxic coordinate, and after a delay of 2 minutes, virus was injected at a rate of 100nl/min. The pipette was left in position for 5 minutes after virus injection to reduce backflow up the injection tract. After pipette removal, the skin was sutured (Ethicon) and sealed with glue (Vetbond). Mice were closely monitored during the recovery phase, and placed in a clean cage on a warmed surface with access to a softened chow diet. Animals were given 2 weeks to recover, and for the virus to express, before being euthanized for perfusion/fixation.

Stereotaxic coordinates: Targeting of dorsal hippocampus (all with respect to Bregma): [AP: -2 | ML: 1.5 | Z: -1.5, -1.3, -1.1 (from pia, 300nl at each site)]. Note, for SLED.NPL deep cortical layers above the hippocampus were imaged (with overflow virus injection into the dorsal hippocampus).

### Perfusions and immunohistochemistry

Mice were terminally anesthetized and transcardially perfused with PBS followed by 4% PFA (Electron Microscopy Sciences), both ice cold. Brains were then postfixed for 2 hours at 4°C and then washed with PBS. Brains were then either sliced on a vibratome (Leica; VT-1000; 60 μm thick) or incubated overnight in 30% sucrose and cut into 40**μ**m sections using a cryostat (Leica Biosystems). For slices that required Gad67 staining the following IHC protocol was followed as previously described^4^. Sections were washed x3 in PBS (10 minutes each) and permeabilized with PBS containing 0.1% Triton-X (Sigma) for 30 minutes at RT. Slices were blocked with PBS containing 3% BSA and 5% normal goat serum (Vector Laboratories) for 1 hour at RT. Primary antibodies were made up in the same blocking buffer and incubated at RT for 24hrs.

Slices were washed 4x with PBS and then incubated with fluorescently conjugated secondary antibodies (Invitrogen, all at 1:500) ON at 4°C. Slices were then washed x4 with PBS and mounted on slides with PermaFluor (Thermo Fisher Scientific). For other antibody combinations the same overall protocol was followed with the following differences. Slices were permeabilized with PBS containing 0.3% Triton-X for 20 minutes. Slices were blocked with PBS containing 5% normal goat serum with the addition of 0.15% Trixon-X. Primary antibodies were incubated in the same blocking buffer but at 4°C overnight. Secondary antibodies were made up in the same blocking buffer and incubated ON at 4°C. In some instances, slices were washed with PBS containing DAPI to label nuclei after the secondary antibody incubation. For SLED.MEv2 the fluorophores were not antibody boosted. For evaluation of SLED.NPL and SLED.ENS the EGFP and mCherry/dsRed was antibody boosted. Slides were imaged on a confocal microscope (as described above), or on an Apotome epifluorescence scope (Zeiss) and analyzed further in Fiji.

### Ferret Cortex AAV injections

An adult female ferret (Mustela putoris furo, Marshall Farms) was used for the virus injection. Anesthesia was induced with ketamine (40 mg/kg) and maintained with isoflurane (1.5 – 3%). Atropine (0.05 mg/kg) was given at the start of the surgery. Burenoprhine (0.01 – 0.03 mg/kg) was administered pre- and post-operatively for analgesia in combination with a subcutaneous injection of lidocaine during the surgery, and post-operative administration of meloxicam (0.1 – 0.2 mg/kg). Animals were maintained at normal body temperature during the surgery using a heating pad. Skin and muscle over primary visual cortex were reflexed, and a small craniotomy was made over the brain region of interest. Virus was then injected through a pulled glass pipette sharpened to a tip angle of about 60 deg. Approximately 1 uL of virus was then injected, distributed across multiple depths at a single site in the craniotomy. After the injection, muscle and skin were closed and the animal was recovered and returned to its home cage. The animal was perfused 3 months after the virus injection.

### Ocular AAV injections

For subretinal injections, AAVs were injected into the subretinal space of 28 day old Prph2^*rds/rds*^ mice (#001979 Jackson Laboratory, Bar Harbor, ME). Briefly, mice were anesthetized by intraperitoneal injection of ketamine (100 mg/kg) and xylazine hydrochloride (20 mg/kg). The pupils were dilated with 1% tropicamide (Alcon, Ft. Worth, TX). The corneas were covered with Healon GV sodium hyaluronate solution (Abbott Medical Optics Inc., Santa Ana, CA) and cover glass to facilitate transpupillary visualization. 1uL of AAV (10^13 viral genomes/mL) were loaded into a 33G needle micro-syringe (Hamilton Company, Reno, NV), then tangentially injected into the subretinal space through the sclera of the mice. A successful injection was verified by direct visualization through the dilated pupil of the recipient under the surgical microscope (Leica, Wetzlar, Germany).

For intravitreal injections, AAVs were injected into the vitreous cavity of 0 day old CD1 pups (Charles River Laboratories, Wilmington, MA). The neonatal animals were anesthetized by placing them on a waterproof surface over crushed ice until the pup was no longer responsive to touch. The eyelids were surgically separated before injecting 1 μL of AAV (10^13 viral genomes/mL) into the vitreous space using a custom 33G sharp needle micro-syringe (Hamilton Company, Reno, NV). The needle was held in place for 10 seconds to avoid outflow before being gently removed.

### Spectral Domain-Optical Coherence Tomography

For *in vivo* retinal imaging, Spectral Domain-OCT images were obtained and analyzed as previously described ^84^. Mice were first anesthetized with ketamine (100 mg/kg) and xylazine hydrochloride (20 mg/kg), followed by dilation with 1% tropicamide and 2.5% phenylephrine. The clarity of the cornea and lens was maintained using GenTeal lubricating eye gel (Novartis Pharmaceuticals, Basel, Switzerland). The mice were secured using a bite bar to a movable stage. The stage was adjusted manually to center the image of the retina at the optic nerve head. Cross-sectional images were generated using 1000 rectangular volume scans using the Envisu OCT system (Leica Microsystems, Wetzlar, Germany). Outer nuclear layer and inner nuclear layer thickness was measured using the linear caliper function in the software by a masked observer using a pre-established uniform grid.

### Electroretinography (ERG)

Full-field flash ERGs were performed as previously described^85^. In brief, mice were dark adapted overnight and anesthetized with ketamine (80 mg/kg) and xylazine hydrochloride (16 mg/kg) prior to recording. The pupils were dilated with 1% tropicamide, 1% cyclopentolate and 2.5% phenylephrine and the corneal surface was anesthetized with 0.5% proparacaine HCl eye drops. For recording retinal electrical responses, stainless-steel wire electrodes were placed on the corneas as the active electrodes, contacting the center of the corneal surface through a thin layer of artificial tear. Needle electrodes were subcutaneously inserted into the cheek and the tail as reference and ground electrodes, respectively. To maintain body temperature during the procedure, the animals were placed on a temperature-controlled heating pad. Using the UTAS Bigshot ERG system (LKC Technologies, Gaithersburg, MD), ERG responses were differentially amplified (0.3–300 Hz), digitized at 1,000 Hz, averaged and stored. The recording epoch was 512 ms, with a 20 ms pre-stimulation baseline. After 7 min of light adaptation, ERGs were obtained to strobe flashes (−0.8 to 1.9 log cd.s/m^2^) superimposed upon a steady 30 cd.s/m^2^ white background. in response to a series of flashes ranging from -0.8 to 1.9 log cd.s/m^2^. The b-wave amplitude was measured from the a-wave trough to the peak of the b-wave.

## Acknowledgements

This work was supported by NIH grants RF1MH123237 (S.B.), R24EY027283 (S.B.), K08EY027093 (F.R.), and R01EY033103 (M.S.S.), a Stein Innovation Award from Research to Prevent Blindness to S.B., an unrestricted departmental grant to the Wilmer Eye Institute from Research to Prevent Blindness awarded to F.R., an NSF NeuroNex grant #1934288 awarded to K.J.N., a Visual Sciences Training grant 2T32EY007143 awarded to C.P.S., a Johns Hopkins Kavli NDI fellowship awarded to J.P.L., and a Johns Hopkins IDIES Seed Fund awarded to J.P.L. We thank the Ross Flow Cytometry Core (JHMI), the Wilmer Microscopy module supported by the EY001765 core grant for flow cytometry, the Single Cell & Transcriptomics Core (JHMI), and the Cleveland Clinic core grant (EY025585). We thank W. Yap, W. Xin, and R. Roth for comments on the manuscript.

## Author contributions

S.B. and J.P.L. conceived and oversaw all aspects of the study. J.P.L., A.M.B., C.P.S., R.P.C.O., V.T., M.Y.G., R.D., L.Z., T.A.B., K.T., J.H., Y.Li and P.J.L. analyzed cellular specificity of SLED vectors. Y.Liu and M.S.S. assisted with subretinal injections. Y.Li and S.S. assisted with human iPSC neuronal culture. D.B., P.P., P.O.K., and T.A.B. assisted with development of calcium indicator SLED vectors. M.Y. and N.P. conducted ERG analysis. B.L. assisted with computational efforts. R.L.H. supervised analysis of SLED.ENS and SLED.NPL vectors. W.B.D., F.R., B.D., K.T. and J.H. carried out studies of oncolytic SLED vectors. K.J.N., J.J.S., and R.V. carried out all ferret studies. J.P.L., A.M.B, C.P.S. and S.B. drafted the manuscript. All authors approved the final manuscript.

