## Supplemental Figures 1-7 for "Cell-specific regulation of gene expression using splicing-dependent frameshifting"

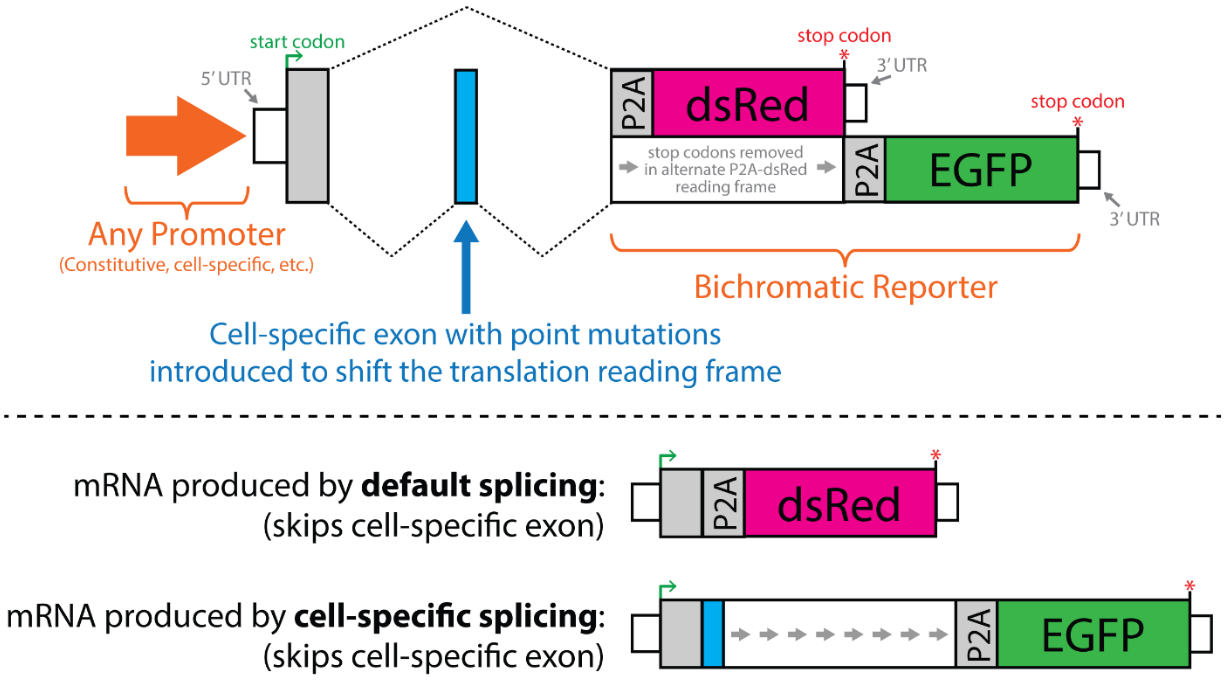


**Supplemental Figure 1.** Detailed schematic of the generic bichromatic reporter used for testing SLED vector specificity. Note that P2A elements are upstream of each fluorescent reporter to avoid interference from N-terminal peptides.

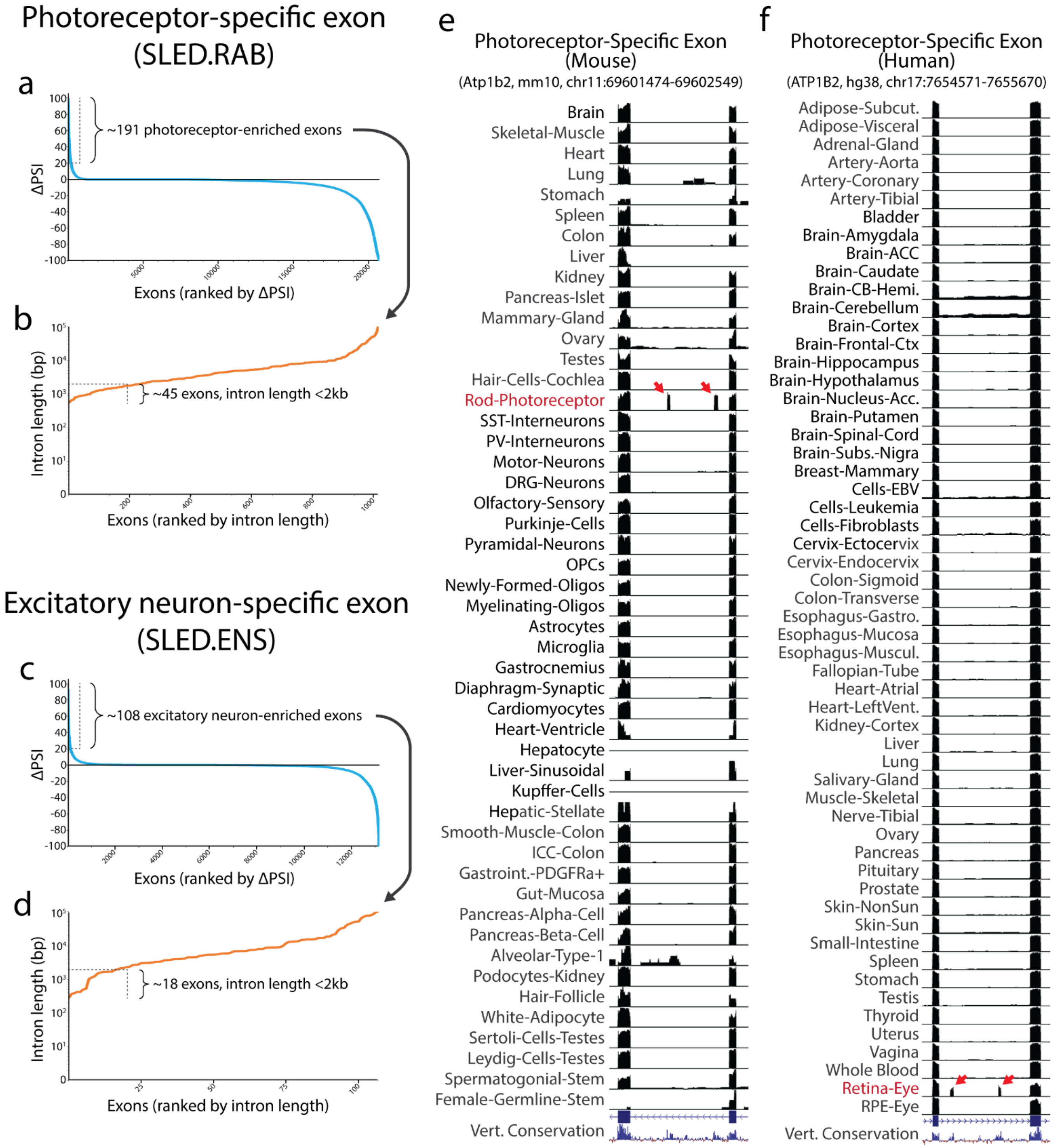

Photoreceptor-specific exon (SLED.RAB)
a
~191 photoreceptor-enriched exons
ΔPSI
Exons (ranked by ΔPSI)
b
Intron length (bp)
~45 exons, intron length <2kb
Exons (ranked by intron length)
Excitatory neuron-specific exon (SLED.ENS)
c
~108 excitatory neuron-enriched exons
ΔPSI
Exons (ranked by ΔPSI)
d
Intron length (bp)
~18 exons, intron length <2kb
Exons (ranked by intron length)
e
Photoreceptor-Specific Exon (Mouse)
(Atp1b2, mm10, chr11:69601474-69602549)
Brain
Skeletal-Muscle
Heart
Lung
Stomach
Spleen
Colon
Liver
Kidney
Pancreas-Islet
Mammary-Gland
Ovary
Testes
Hair-Cells-Cochlea
Rod-Photoreceptor
SST-Interneurons
PV-Interneurons
Motor-Neurons
DRG-Neurons
Olfactory-Sensory
Purkinje-Cells
Pyramidal-Neurons
OPCs
Newly-Formed-Oligos
Myelinating-Oligos
Astrocytes
Microglia
Gastrocnemius
Diaphragm-Synaptic
Cardiomyocytes
Heart-Ventricle
Hepatocyte
Liver-Sinusoidal
Kupffer-Cells
Hepatic-Stellate
Smooth-Muscle-Colon
ICC-Colon
Gastroint.-PDGFRa+
Gut-Mucosa
Pancreas-Alpha-Cell
Pancreas-Beta-Cell
Alveolar-Type-1
Podocytes-Kidney
Hair-Follicle
White-Adipocyte
Sertoli-Cells-Testes
Leydig-Cells-Testes
Spermatogonial-Stem
Female-Germline-Stem
Vert. Conservation
f
Photoreceptor-Specific Exon (Human)
(ATP1B2, hg38, chr17:7654571-7655670)
Adipose-Subcut.
Adipose-Visceral
Adrenal-Gland
Artery-Aorta
Artery-Coronary
Artery-Tibial
Bladder
Brain-Amygdala
Brain-ACC
Brain-Caudate
Brain-CB-Hemi.
Brain-Cerebellum
Brain-Cortex
Brain-Frontal-Ctx
Brain-Hippocampus
Brain-Hypothalamus
Brain-Nucleus-Acc.
Brain-Putamen
Brain-Spinal-Cord
Brain-Subs.-Nigra
Breast-Mammary
Cells-EBV
Cells-Leukemia
Cells-Fibroblasts
Cervix-Ectocervix
Cervix-Endocervix
Colon-Sigmoid
Colon-Transverse
Esophagus-Gastro.
Esophagus-Mucosa
Esophagus-Muscul.
Fallopian-Tube
Heart-Atrial
Heart-LeftVent.
Kidney-Cortex
Liver
Lung
Salivary-Gland
Muscle-Skeletal
Nerve-Tibial
Ovary
Pancreas
Pituitary
Prostate
Skin-NonSun
Skin-Sun
Small-Intestine
Spleen
Stomach
Testis
Thyroid
Uterus
Vagina
Whole Blood
Retina-Eye
RPE-Eye
Vert. Conservation

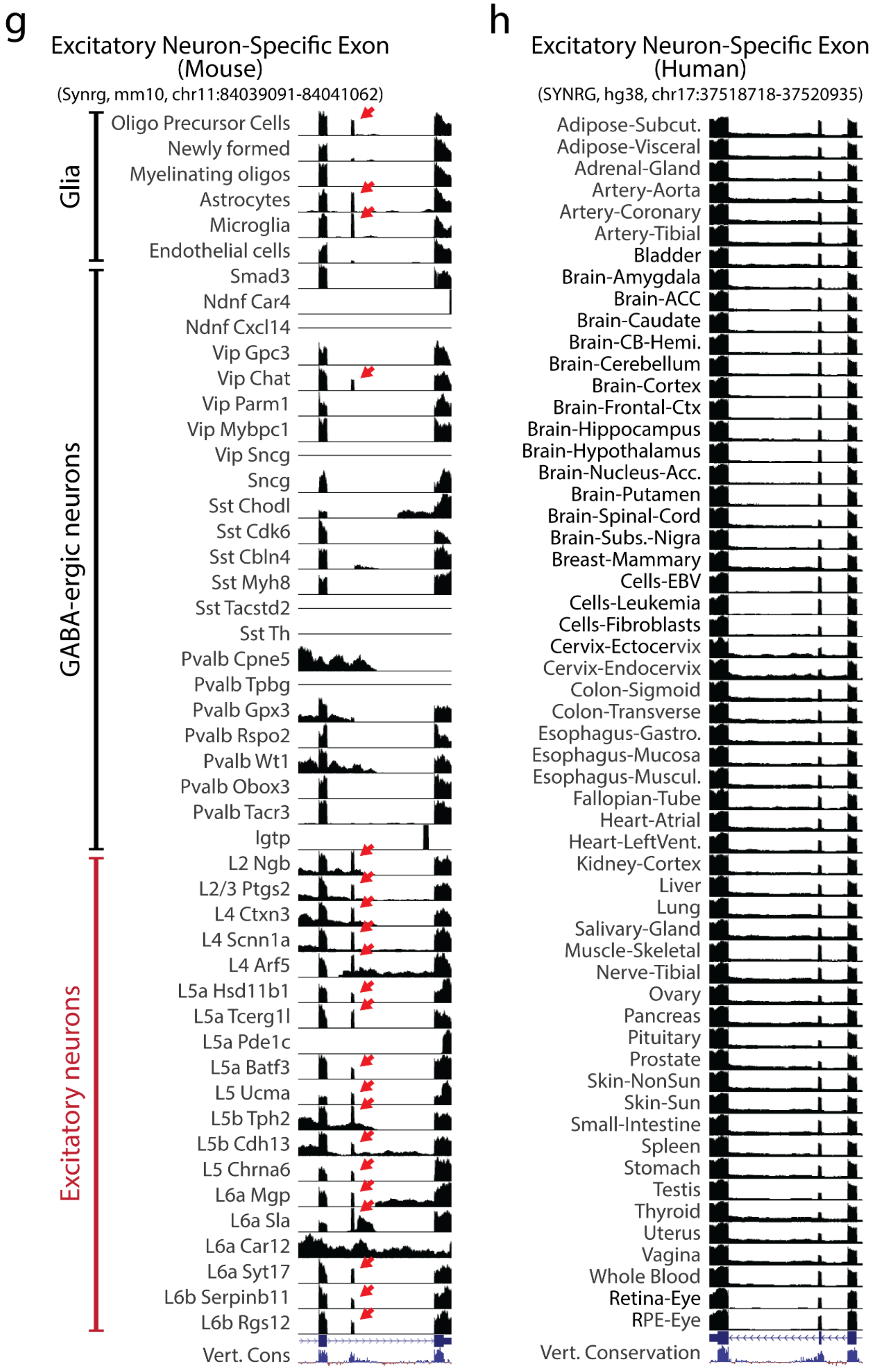
g
Excitatory Neuron-Specific Exon (Mouse)
(Synrg, mm10, chr11:84039091-84041062)
Glia
Oligo Precursor Cells
Newly formed
Myelinating oligos
Astrocytes
Microglia
Endothelial cells
GABA-ergic neurons
Smad3
Ndnf Car4
Ndnf Cxcl14
Vip Gpc3
Vip Chat
Vip Parm1
Vip Mybpc1
Vip Sncg
Sncg
Sst Chodl
Sst Cdk6
Sst Cbln4
Sst Myh8
Sst Tacstd2
Sst Th
Pvalb Cpne5
Pvalb Tpbg
Pvalb Gpx3
Pvalb Rspo2
Pvalb Wt1
Pvalb Obox3
Pvalb Tacr3
Igtp
Excitatory neurons
L2 Ngb
L2/3 Ptgs2
L4 Ctxn3
L4 Scnn1a
L4 Arf5
L5a Hsd11b1
L5a Tcerg1l
L5a Pde1c
L5a Batf3
L5 Ucma
L5b Tph2
L5b Cdh13
L5 Chrna6
L6a Mgp
L6a Sla
L6a Car12
L6a Syt17
L6b Serpinb11
L6b Rgs12
Vert. Cons
h
Excitatory Neuron-Specific Exon (Human)
(SYNRG, hg38, chr17:37518718-37520935)
Adipose-Subcut.
Adipose-Visceral
Adrenal-Gland
Artery-Aorta
Artery-Coronary
Artery-Tibial
Bladder
Brain-Amygdala
Brain-ACC
Brain-Caudate
Brain-CB-Hemi.
Brain-Cerebellum
Brain-Cortex
Brain-Frontal-Ctx
Brain-Hippocampus
Brain-Hypothalamus
Brain-Nucleus-Acc.
Brain-Putamen
Brain-Spinal-Cord
Brain-Subs.-Nigra
Breast-Mammary
Cells-EBV
Cells-Leukemia
Cells-Fibroblasts
Cervix-Ectocervix
Cervix-Endocervix
Colon-Sigmoid
Colon-Transverse
Esophagus-Gastro.
Esophagus-Mucosa
Esophagus-Muscul.
Fallopian-Tube
Heart-Atrial
Heart-LeftVent.
Kidney-Cortex
Liver
Lung
Salivary-Gland
Muscle-Skeletal
Nerve-Tibial
Ovary
Pancreas
Pituitary
Prostate
Skin-NonSun
Skin-Sun
Small-Intestine
Spleen
Stomach
Testis
Thyroid
Uterus
Vagina
Whole Blood
Retina-Eye
RPE-Eye
Vert. Conservation

i

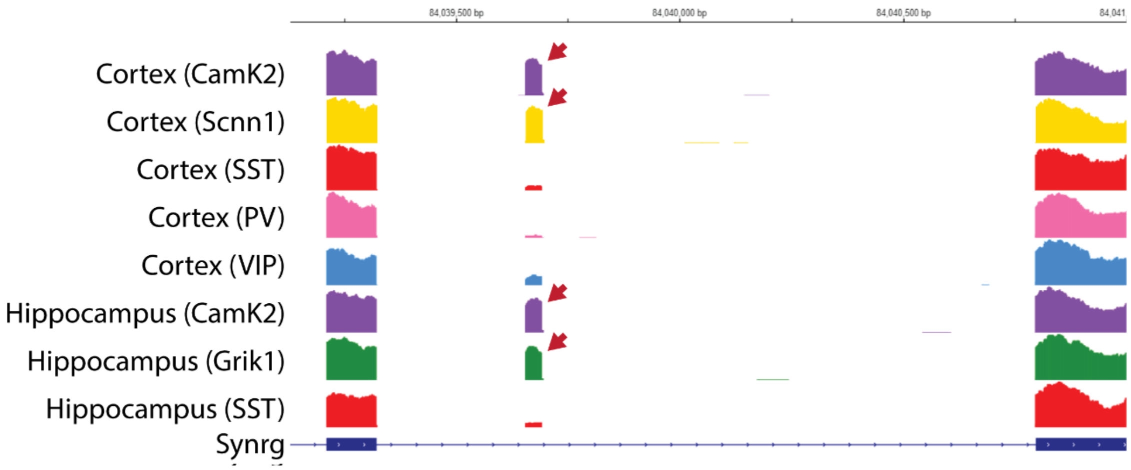


**Supplemental Figure 2.** Using the computational resource ASCOT to identify photoreceptor-specific and excitatory neuron-specific exon candidates for SLED vector design. For the photoreceptor-specific exon, we identified ~191 photoreceptor-enriched alternative exons (**a**), of which ~41 had intronic lengths of <2 kb (**b**). A photoreceptor-enriched exon in the gene *Atp1b2* was selected for characterization. For the excitatory neuron-specific exon, we identified ~108 exons enriched in excitatory neurons compared to GABAergic neurons (**c**), of which ~18 had intronic lengths of <2 kb (**d**). Using ASCOT, an excitatory neuron-enriched exon in the gene *Synrg* was chosen for characterization. UCSC track views of the *Atp1b2* exon for mouse (**e**) and human (**f**) and UCSC track views of the *Synrg* exon in mouse (**g**) and human (**h**). Datasets in (g) are from V1 cortex (Tasic et al., Nat. Neuro. 2016). Note that the *Synrg* is only selective when expressed under a neuron-specific promoter like hSyn, as the exon is spliced in other non-neuronal tissues and cell types such as OPCs, astrocytes, and microglia. Importantly, while the *Synrg* exon was determined to be the best candidate for excitatory neuron SLED, it does exhibit expression in a limited subset of non-excitatory neurons, most notably the Chat$^{+}$ subtype of VIP neurons in V1 cortex splice-in the *Synrg* exon. No clear cell specificity is observed in human GTEx data, as these datasets were sequenced from whole tissues (**h**). In a recent set of high depth RNA-Seq datasets generated from cortex and hippocampal neurons using RiboTRAP (Furlanis*, Lisa Traunmüller* et al., Nat. Neuro. 2019), the *Synrg* exon exhibits high PSI in excitatory neurons and low PSI in inhibitory neurons (**i**).

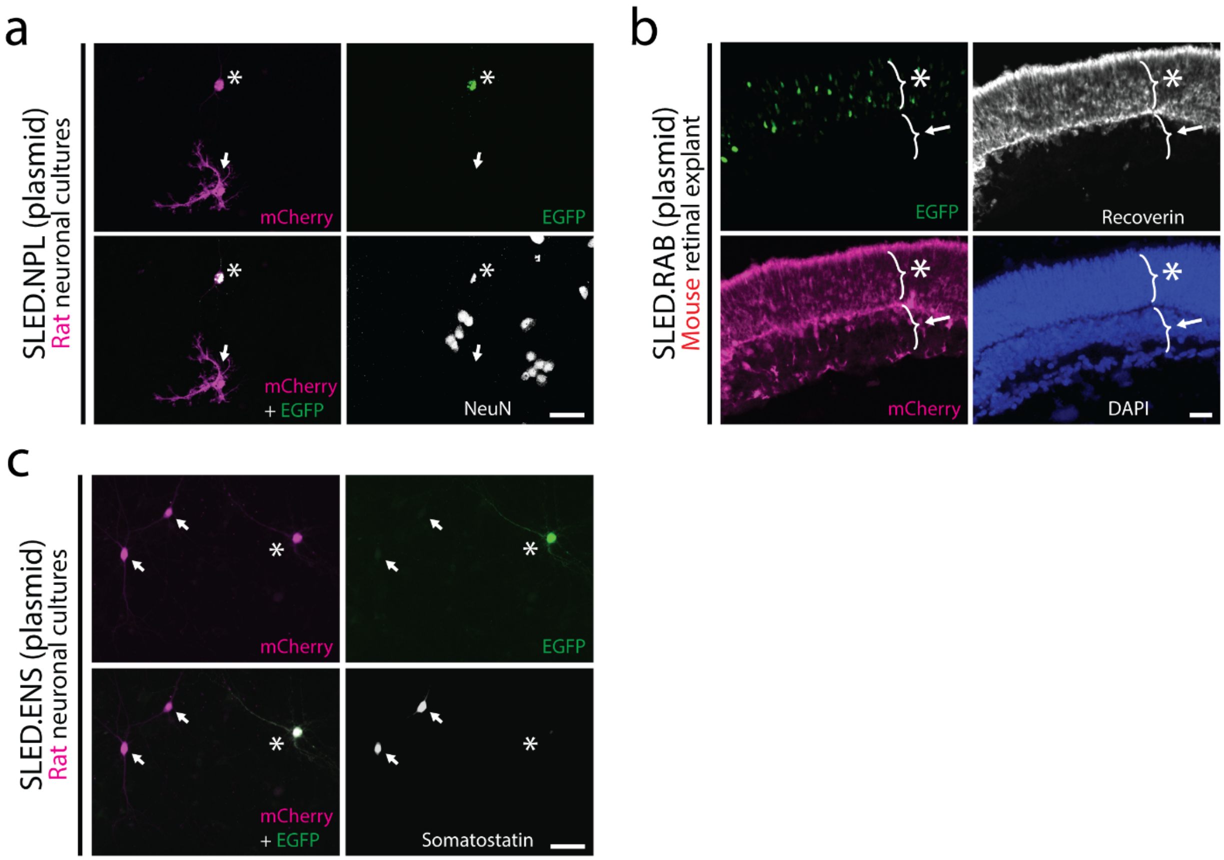


**Supplemental Figure 3.** Proof-of-concept testing of SLED.NPL, SLED.RAB, and SLED.ENS using *in vitro* transfection and electroporation. (**a**) SLED.NPL was transfected into primary rat neuronal cultures and we observed selective expression of EGFP in neurons, while the default splicing-driven dsRed was observed in transfected neurons and glia (asterisk = neuron, arrow = glia). (**b**) SLED.RAB was electroporated into P0 mouse retinas and we observed that while expression of dsRed was present in all postnatally-generated cell types, EGFP expression was restricted to photoreceptors (asterisk = ONL, arrow = INL). (**c**) Transfection of the SLED.ENS constructs into primary rat hippocampal cultures resulted in exclusion of EGFP in somatostatin-positive inhibitory neurons. Splicing-in of the SLED-ENS Synrg exon is coupled to the EGFP reading frame. Asterisk = putative excitatory neuron, arrows = $SST^{+}$ GABAergic interneurons. Scale bars = 50μm.

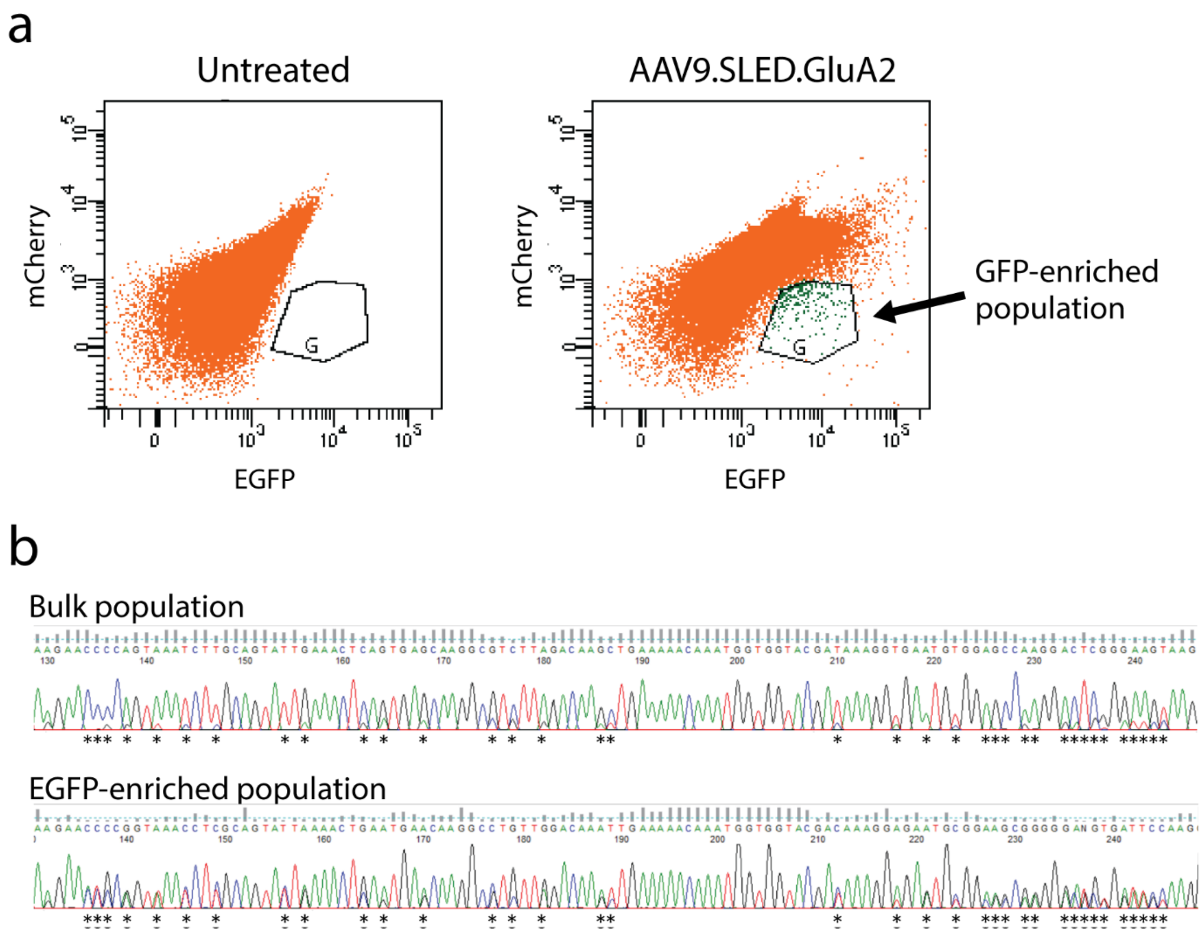


**Supplemental Figure 4.** (**a**) FACS isolation of an mCherry$^{low}$ and EGFP$^{high}$ population from AAV9.SLED.GluA2 transduced neurons (rat primary neuronal culture). (**b**) Sanger sequencing of AAV.SLED.GluA2 RTPCR products generated from the bulk and GFP-enriched populations. Flip and flop exons are nearly identical in sequence. However, at basepair positions that diverge (asterisks), chromatogram signals associated with the flop variant are observed at a much higher frequency in the GFP-enriched population when compared to bulk.

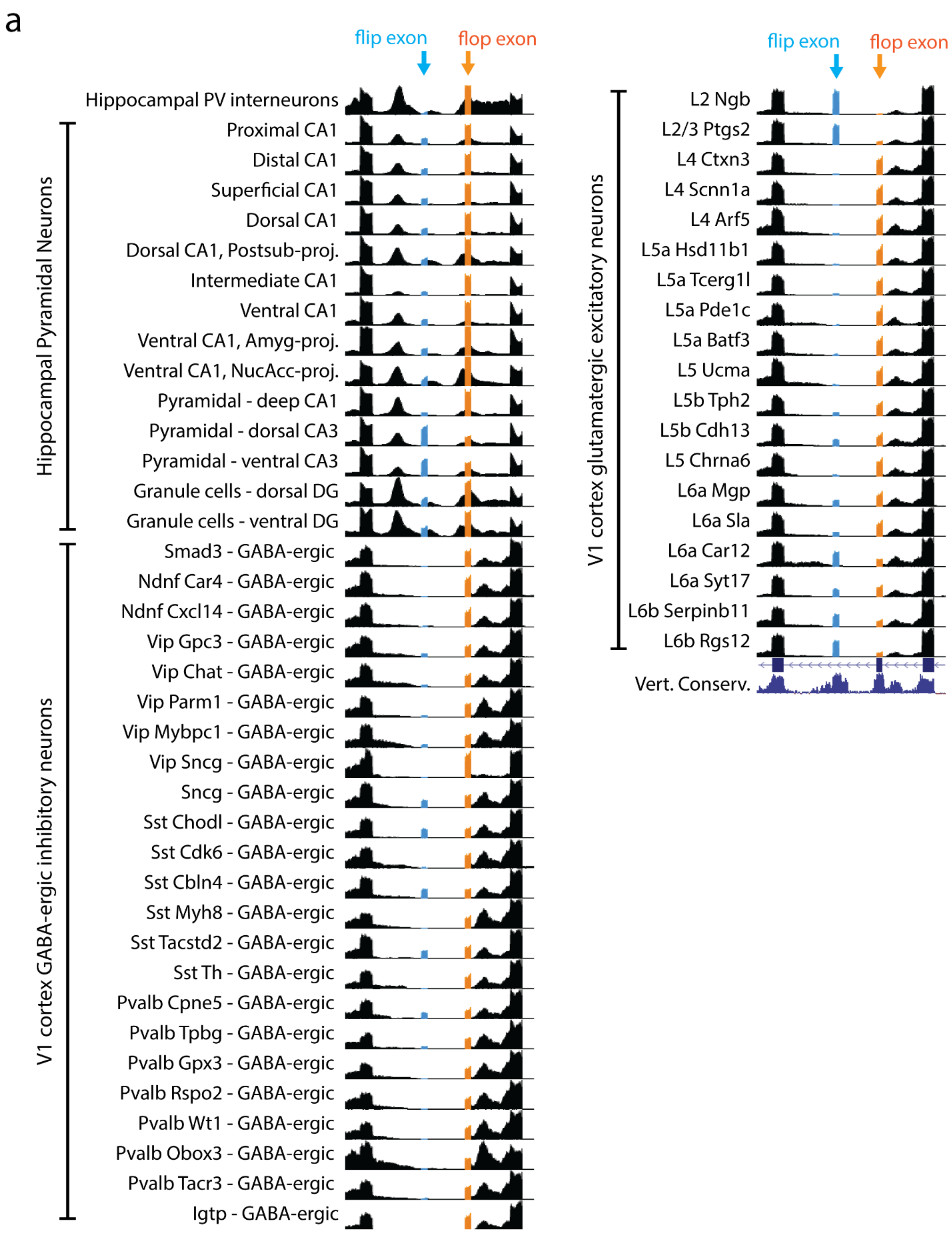
a
flip exon
flop exon
Hippocampal PV interneurons
Proximal CA1
Distal CA1
Superficial CA1
Dorsal CA1
Dorsal CA1, Postsub-proj.
Intermediate CA1
Ventral CA1
Ventral CA1, Amyg-proj.
Ventral CA1, NucAcc-proj.
Pyramidal - deep CA1
Pyramidal - dorsal CA3
Pyramidal - ventral CA3
Granule cells - dorsal DG
Granule cells - ventral DG
Hippocampal Pyramidal Neurons
Smad3 - GABA-ergic
Ndnf Car4 - GABA-ergic
Ndnf Cxcl14 - GABA-ergic
Vip Gpc3 - GABA-ergic
Vip Chat - GABA-ergic
Vip Parm1 - GABA-ergic
Vip Mybpc1 - GABA-ergic
Vip Sncg - GABA-ergic
Sncg - GABA-ergic
Sst Chodl - GABA-ergic
Sst Cdk6 - GABA-ergic
Sst Cbln4 - GABA-ergic
Sst Myh8 - GABA-ergic
Sst Tacstd2 - GABA-ergic
Sst Th - GABA-ergic
Pvalb Cpne5 - GABA-ergic
Pvalb Tpbg - GABA-ergic
Pvalb Gpx3 - GABA-ergic
Pvalb Rspo2 - GABA-ergic
Pvalb Wt1 - GABA-ergic
Pvalb Obox3 - GABA-ergic
Pvalb Tacr3 - GABA-ergic
Igtp - GABA-ergic
V1 cortex GABA-ergic inhibitory neurons
flip exon
flop exon
L2 Ngb
L2/3 Ptgs2
L4 Ctxn3
L4 Scnn1a
L4 Arf5
L5a Hsd11b1
L5a Tcerg1l
L5a Pde1c
L5a Batf3
L5 Ucma
L5b Tph2
L5b Cdh13
L5 Chrna6
L6a Mgp
L6a Sla
L6a Car12
L6a Syt17
L6b Serpinb11
L6b Rgs12
V1 cortex glutamatergic excitatory neurons
Vert. Conserv.

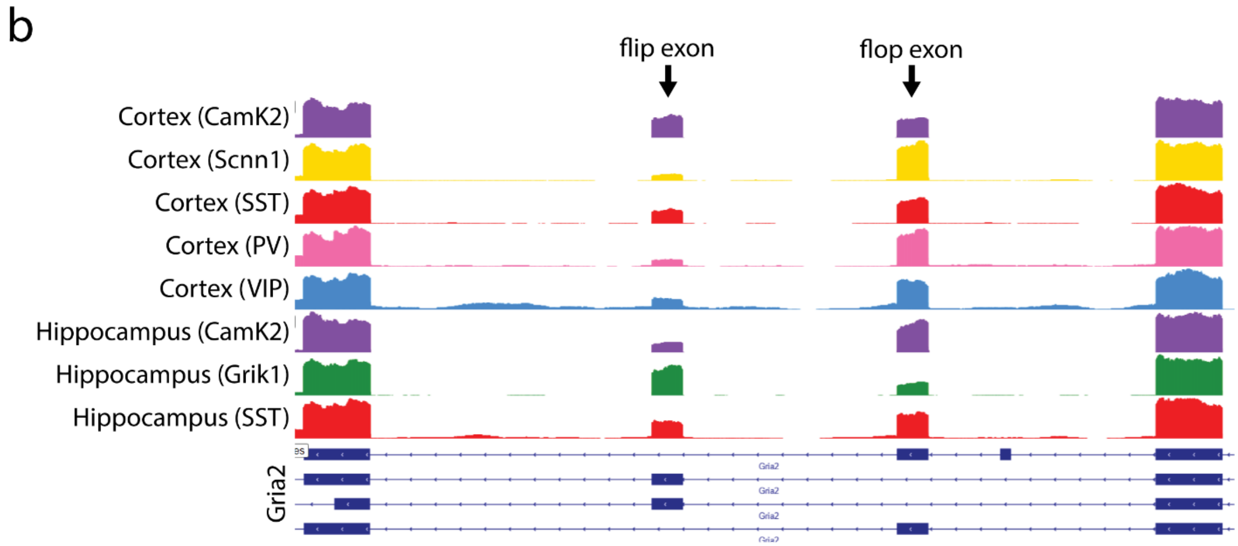


**Supplemental Figure 5.** UCSC track views of GluA2 flip/flop splicing from publicly archived single-cell SMART-Seq datasets in mouse V1 cortex (**a**) (Tasic et al., Nat. Neuro. 2016) and bulk neuronal data using RiboTRAP mice (**b**) (Furlanis*, Lisa Traunmüller* et al., Nat. Neuro. 2019). Glutamatergic neurons typically show higher expression of the flop exon while GABAergic neurons predominantly express the flip exon, but there is still extensive variability in GluA2 flip/flop ratios between cell types in cortex and hippocampus.

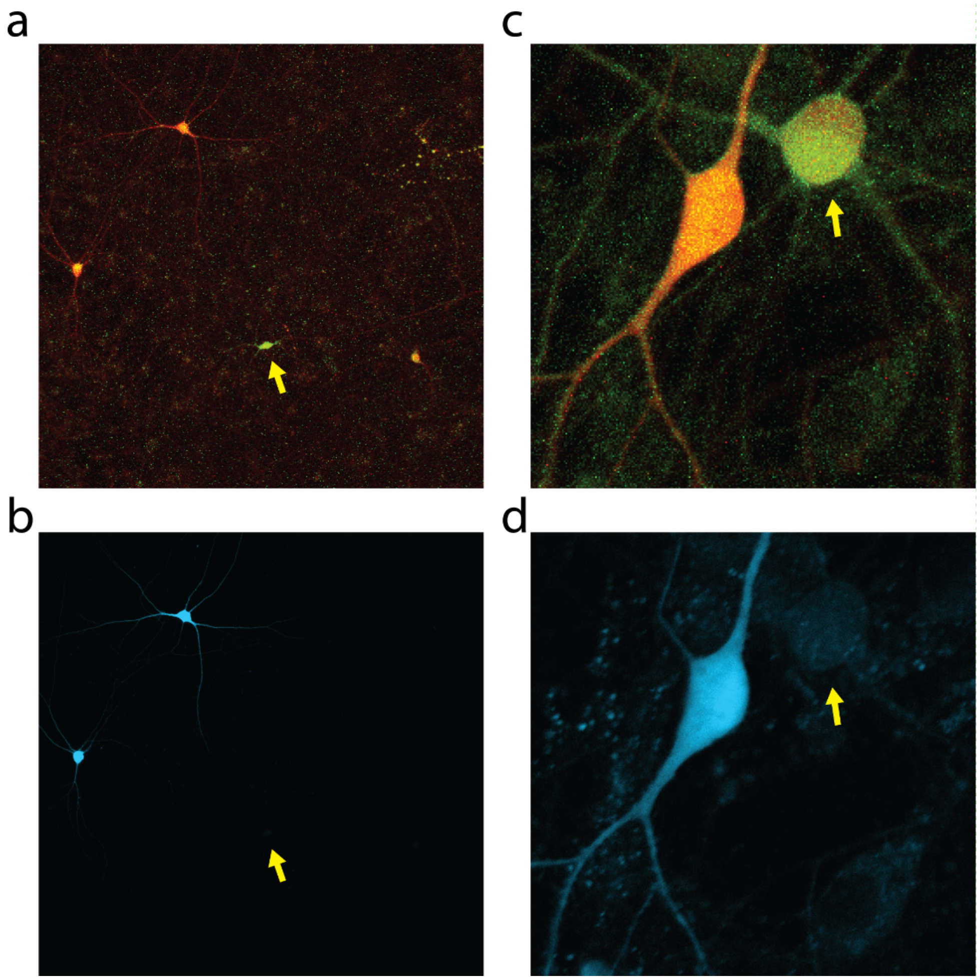


**Supplemental Figure 6.** (**a**) Maximum intensity projection of RGECO1a and GCaMP7b composite images used to generate Supplemental Video 1. (**b**) mDlx-Azurite expression indicates inhibitory neurons while the mDlx-Azurite negative cell is an excitatory neuron (arrow). (**c**) Maximum intensity projection of RGECO1a and GCaMP7b composite images used to generate Supplemental Video 2. (**d**) mDlx-Azurite expression indicates inhibitory neuron while mDlx-Azurite negative cell is an excitatory neuron (arrow).

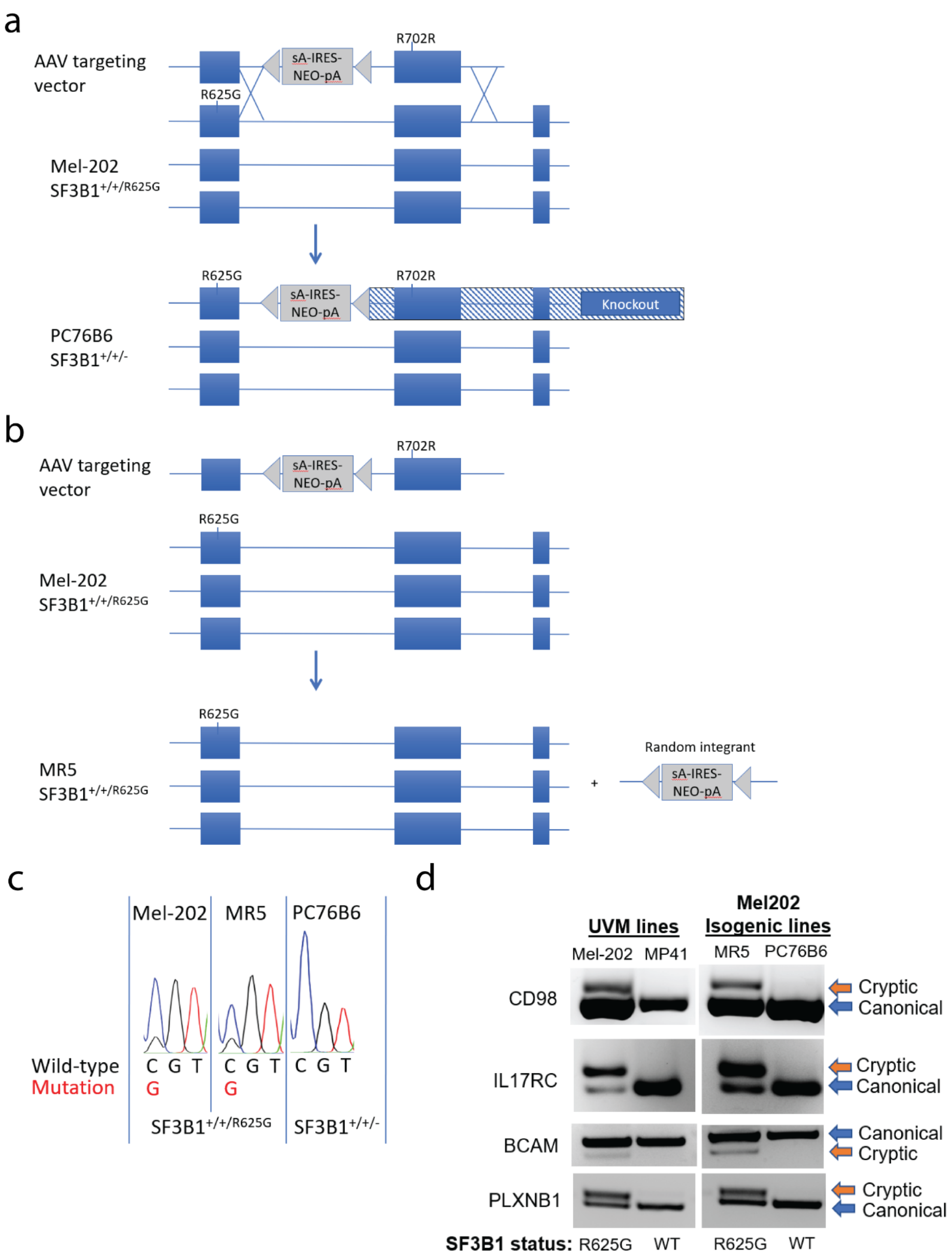
a
AAV targeting vector
sA-IRES-NEO-pA
R702R
R625G
Mel-202 SF3B1+/+/R625G
PC76B6 SF3B1+/+/-
Knockout
b
AAV targeting vector
sA-IRES-NEO-pA
R702R
R625G
Mel-202 SF3B1+/+/R625G
MR5 SF3B1+/+/R625G
Random integrant
c
Mel-202
MR5
PC76B6
Wild-type
Mutation
C G T
G
SF3B1+/+/R625G
SF3B1+/+/-
d
UVM lines
Mel202 Isogenic lines
Mel-202
MP41
MR5
PC76B6
CD98
IL17RC
BCAM
PLXNB1
Cryptic
Canonical
SF3B1 status: R625G WT R625G WT

**Supplemental Figure 7.** Engineering and validation of isogenic Mel-202 cells. (**a**) Targeting event creating PC76B6 cells, with a functional knockout of *SF3B1*$^{R625G}$. (**b**) Gene targeting control clone MR5 exposed to AAV and single-cell isolation, but which remains mutant, showing random integration of NEO cassette. (**c**) Sanger sequencing of cDNA for the *SF3B1* gene, showing disappearance of *SF3B1*$^{R625G}$ expression in PC76B6 cells. (**d**) Validation by RT-PCR of reversion of *SF3B1*$^{R625G}$ cryptic splicing events in PC76B6 cells.

**Supplemental Video 1. CaRPv1_10x.mp4**
Video of primary rat neuronal cultures transfected with SLED.CaRPv1 and imaged using a 10x objective at 2Hz (framerate = 7fps). Images were normalized and processed according to the protocol listed in the Methods section.

**Supplemental Video 2. CaRPv1_20x.mp4**
Video of primary rat neuronal cultures transfected with SLED.CaRPv1 (bicuculline treated) and imaged using a 20x objective at 4Hz (framerate = 30fps). Although transfection of primary rat cultures leads is sparse, occasionally transfected neurons are close enough in proximity for simultaneous imaging at 20x magnification. Images were normalized and processed according to the protocol listed in the Methods section.
